# Modulation of spatial cue processing across the lifespan: a geometric polarization of space restores allocentric navigation strategies in children and older adults

**DOI:** 10.1101/2020.02.12.945808

**Authors:** Marcia Bécu, Denis Sheynikhovich, Stephen Ramanoël, Guillaume Tatur, Anthony Ozier-Lafontaine, José-Alain Sahel, Angelo Arleo

**Author notes:** corresponding author Contact information:, postal address: 17, rue Moreau, 75012, Paris, France.

## Abstract

The impact of development and healthy aging on spatial cognition has been traditionally attributed to a difficulty in using allocentric strategies and a preference for egocentric ones. An alternative possibility, suggested by our previous works, is that this preference is actually conditioned by the spatial cues (e.g. geometric of landmark cues) present in the environment rather than a strategic choice per se. We tested this prediction by having 79 subjects (children, young and older adults) navigating a Y-maze composed either of landmarks or geometric cues, with an immersive head-mounted display that allows us to record both head and eye movements. Our results show that when the performance is based on landmarks solely, children and older adults exhibit a deficit in using allocentric strategies when compared to young adults. Hence, an inverted U-profile of allocentric strategies was observed across the lifespan. This was not due to a default of attention to the landmarks, as evidenced by analysis of gaze dynamics. When geometric were provided, however, older adults and children used allocentric strategies in the same proportion as young adults. They were, in addition, as efficient and quick to implement the strategy. We thus propose a reinterpretation of the previous data in the literature, whereby reference to geometric cues is the default mode for spatial representations, which is immune to age, whereas spatial representations fail to be anchored on landmarks early in development and later in aging. This new interpretation has the potential to reunify several data from the literature, ranging from spatial cues processing to strategy preference, and including other spatial skills like path integration and route learning.

---

Spatial information about location can be represented according to two reference frames. In an egocentric frame of reference, locations are represented relative to the subject’s body, whereas in an allocentric frame of reference, locations are represented relative to external environmental elements^1^. The use of allocentric and egocentric strategies is highly influenced by specific situations the navigator is facing^2^ but also by the navigator’s individual characteristics. Among the later, age critically influence the way people navigate in space^3^.

Developmental and aging data seem to suggest an inverted U-profile of allocentric processing with age. In young children, spatial navigation seems to be preferentially guided by egocentric representations, although some form of allocentric coding can be present early in development^4,5^. For instance, by having children aged between 5 and 10 navigating a multiple-armed maze surrounded by distal landmarks, Bullens et al. (2010) showed that a majority of children used an sequential egocentric strategy spontaneously^6^. The use of allocentric strategies emerged gradually later on during development, from 7 to 10 years of age. At the other end of the curve, converging evidence supports a decreased use of allocentric strategies (and a preference for egocentric ones) in older subjects, when compared to young ones^7–12^. For instance, Rodgers et al. (2012) used a paradigm allowing to dissociate allocentric and egocentric strategies in a 3-armed maze (so-called Y-maze)^12^. After having learned to position of a goal starting from one arm of the maze, the subject was positioned in the second arm of the maze and asked to return to the goal position. The subject’s strategy was classified as egocentric if the subject made the same turn as during the learning phase and allocentric if he or she moved to the correct goal location. Results showed that older adults were more likely in this situation to use an egocentric strategy, in comparison to young adults. These age-related shifts in strategy use have been interpreted in relation to the slow maturation (in development) and early deterioration (in aging) of the brain areas involved in spatial navigation (e.g. the hippocampus or frontal areas^13–17^).

An alternative hypothesis has been proposed recently. In a study in ecological conditions, Bécu et al. (2019) showed that healthy aging was associated with an increased preference for geometric spatial cues^18^. Whereas young adults relied on landmark cues, older adults preferentially used geometric cues to reorient in space, at the detriment of the landmarks. This preference for geometry has been linked in older adults to a difficulty in either take perspective or code locations relative to landmarks. These results potentially challenge the classical view for a specific deficit of allocentric strategy in aging, given that standard paradigms that tested navigational strategies are exclusively dependent on the capacity of the subjects to use distal landmarks^8–12^. Furthermore, geometric cues in these paradigms were always unpolarized and could thus not be used for orientation. It is thus possible that age-related navigation difficulties, previously explained by a specific deficit of allocentric strategies, are actually linked to a difficulty in processing landmarks. If this interpretation is true, providing geometric information could potentiate the use of allocentric strategies both in older adults and children, in which a preference for geometric cues has also been shown^19^.

## Results

To test this hypothesis, we adapted the Y-maze paradigm by introducing a geometric polarization to the maze to test the strategy use in a sample of seventy-nine subjects (29 children, 22 young adults and 28 older adults). We used an immersive virtual reality head-mounted display in order for the participant to experience proprioceptive and vestibular inputs while navigating freely in the virtual environment. By recording eye movements while the subject navigated, we sought to unveil potential gaze-related processes that could explain age-related difficulty in using landmarks and to get a deeper insight into the nature of the processes involved during spatial navigation. The participants of this study were randomly assigned to the landmark or the geometry versions. In the landmark version (fig. 1a), the maze had equiangular arms separated by 120° and it was surrounded by three distal landmarks. In the geometry version (fig. 1b), two arms were closer to each other (50°), thus providing a geometric polarization to the maze. The two versions had an equivalent level of difficulty, given that basic measures of spatial learning between the two versions were equivalent in young adults (see supp. results 1 and supp. fig. 1). During the “learning phase”, the subjects had to learn to position of an invisible goal that triggered a rewarding sound (dashed area on fig. 1ab), starting from the same position (position A on fig. 1ab). After having reached the goal position directly for 4 consecutive trials, the “testing phase” started. It consisted of six trials in which the starting position was changed, unknown to the subject, in a pseudo-random manner: three “probe trials” started from B, three “control trials” started from A (see Methods and supp. fig. 2 for a view experienced by the subjects in each version). The probe trials allowed us to test the reliance on external environmental information to code for the goal position. Indeed, if the subject used the distal landmarks (in the landmark version) or the geometric polarization (in the geometry version) to navigate to the actual goal position (C), he/she be using an allocentric strategy for that particular probe trial.

**Figure 1.**
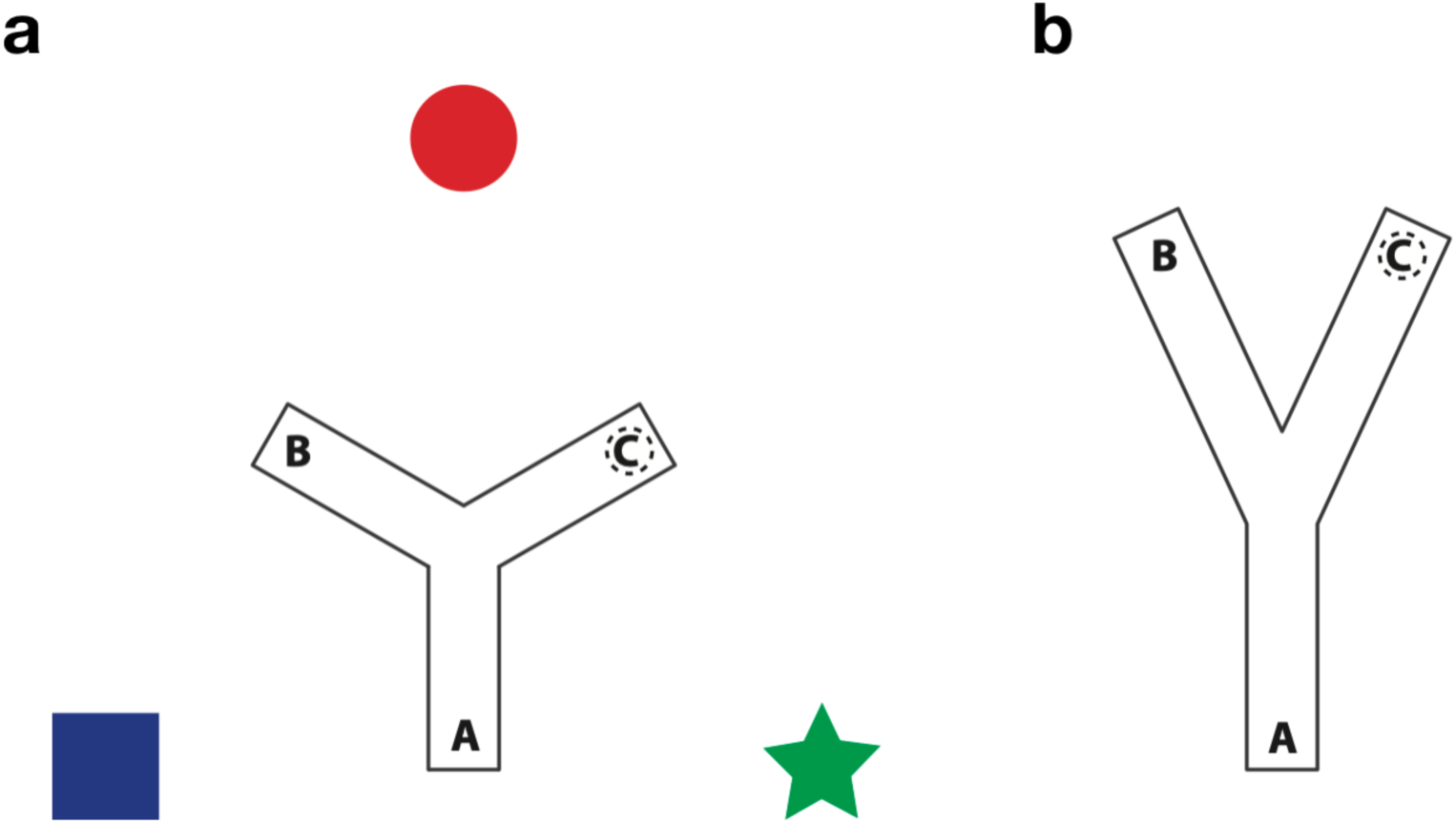
Two versions of the Y-maze. In the landmark version (a), the Y-maze had equiangular arms separated by 120°. Three distal landmarks were placed outside of the maze. In the geometry version (b), an anisotropic arrangement of the 3 arms (50°/155°/155°) provided the geometric polarization. No distal landmarks were present in this condition. Each subject was randomly assigned to the landmark or the geometric version and the experimental procedure was the same for the two versions. After calibrating the eye-tracking device, the subject explored the environment during 3 trials of 1min each, starting from A, B and C (“exploration phase”). Then, the “learning phase” consisted of several trials where the subject had to learn to navigate as directly as possible to the goal position (dashed area C) while always starting from position A. A rewarding signal was automatically displayed when the subject entered the goal area. After four consecutive successful trials, the “testing phase” consisted of 6 trials where the starting position of the subject was changed, unknown to him/her, in a pseudo-random manner: three trials (termed “probe trials”) started from B and 3 trials (termed “control trials”) started from A. No rewarding sound was provided during this phase. During the probe trials, if the subject used the distal landmarks or the geometric polarization to navigate to the actual goal position (C), he is using an allocentric strategy for that particular probe trial. Otherwise, if he keeps on turning right to position A, the subject is using an egocentric strategy, representing the goal position relative to his/her own body position. Participants were disoriented before each trial of the experiment.

Figure 2 shows the proportion of time participants choose an allocentric strategy on the three probe trials. There was some intra-subject variability in the choices made, with subjects either choosing the allocentric option always or only on some of the trials. In the landmark version, we observed an inverted U-profile of allocentric strategy use related to age, with children and older adults being significantly less likely to use an allocentric strategy in this version, relative to young adults (Fisher’s exact probability test in children: p<0.01, ϕ=0.56, oddsratio: 18; in older adults: p<0.01, ϕ=0.53, oddsratio: 16.5). In the geometry version, there was no age difference in the observed proportion of allocentric choices (children: p=0.28, ϕ=0.27; older adults: p=0.22, ϕ=0.32, compared to young adults), with a majority of subjects in each age group using an allocentric strategy (see also supp. fig. 3). Together, these data support the fact that the traditional observation, whereby older adults and children exhibit a specific deficit in the use of allocentric strategies, to the benefit of egocentric ones, was actually erroneous. Indeed, when geometric cues are provided by the environment, children and older adults are just as efficient as young adults to use flexible and more complex allocentric strategies. These results suggest that the impairment previously observed across the lifespan is rather linked to a specific deficit in using landmarks to represent spatial locations.

**Figure 2.**
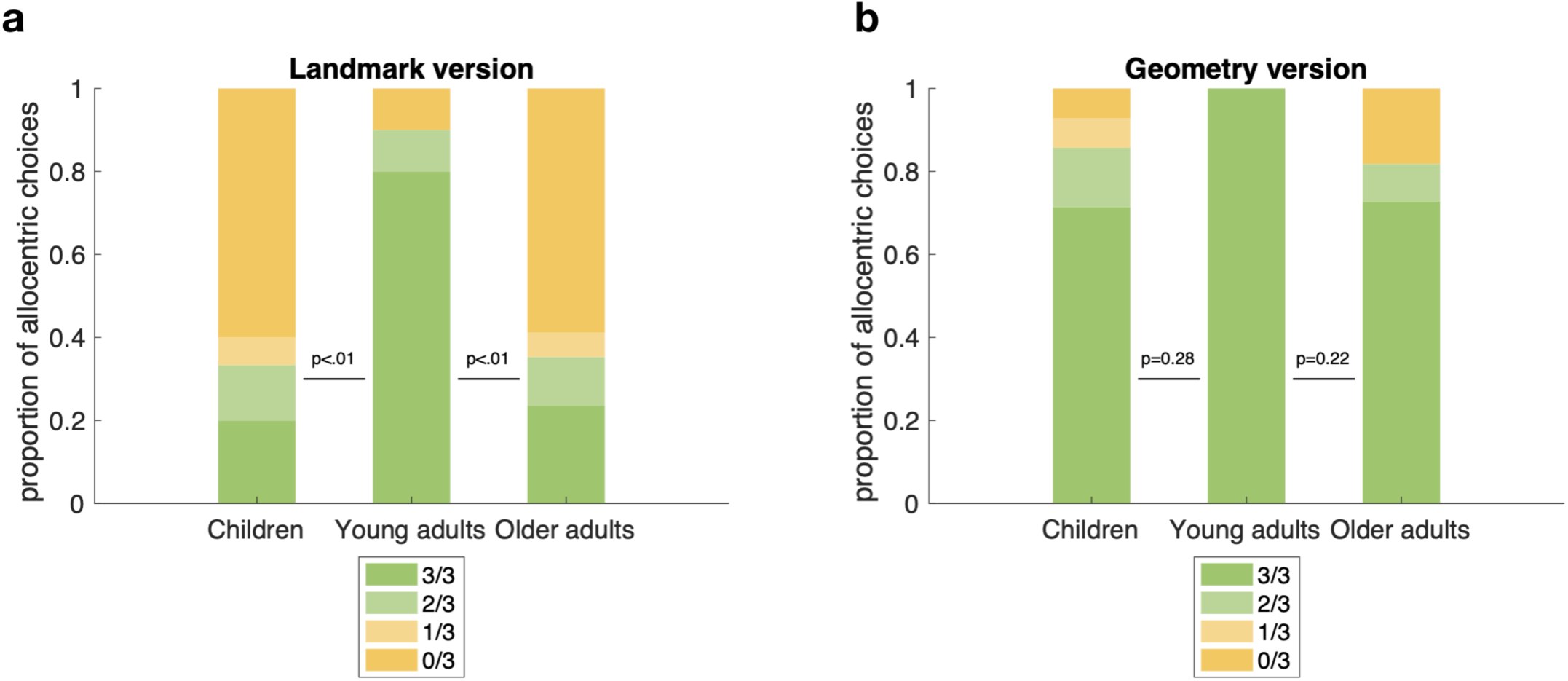
Proportion of allocentric choices in the three probe trials in the landmark (a) and the geometry (b) versions, for the three age groups. In the landmark version (n=42), we observe an inverted U-profile of allocentric strategy use related to age, with children and older adults being significantly less likely to use an allocentric strategy in this version, relative to young adults. We observed no age difference in the geometry version (n=37), supporting the fact that children and older adults depend on the presence of geometric cues to be able to use allocentric strategies as young adults. P-values on the graph indicate Fisher’s exact probability test for two categories: majority of allocentric choices (green color: 3/3 and 2/3) and minority of allocentric choices (yellow color: 1/3 and 0/3). Among the later category, two subjects returned to the starting position (i.e. area B) in the landmark version.

We next wanted to compare how people explored and learned the two environments in order to understand why children and older participants are better at using allocentric strategies in the presence of geometric cues. We estimated several navigation variables based on the trajectory employed by our participants during the learning phase and searched for age differences across the two versions of the maze. Overall, we found that age differences were significant in the landmark version but not in the geometry version, indicating that learning a simple spatial rule (turn right at the intersection) in a maze composed of distal landmarks solely was already more difficult for older adults and children. Indeed, in the landmark version, older adults and children required a higher number of trials in order to reach the learning criterion (i.e. 4 consecutive trials where the subject navigates directly to the goal position, Wilcoxon rank sum test in children vs. young adults: U=226.5,p=0.065; in older adults vs. young adults: U=287.5, p<0.05) whereas in the geometry version the three age groups had a nearly optimal performance, with a median number of trial needed of 4-5 trials (minimum observable: 4, children vs. young adults: U=213.5,p=0.19; older adults vs. young adults: U=141.5,p=0.55, fig. 3a). We then used one-way ANOVA (age factor with 3 modalities) on navigation variables averaged on the first four trials of the learning phase, which were common to all subjects. Age differences were significant in the landmark version for all considered navigation variables (fig. 3b-e and supp. fig. 4, see also supp. fig. 5 for scatter plots) and multiple comparisons of these data indicate that this main effect was due, most of time, to a difference between young adults and the two other groups (see. supp. table 2 for exceptions). In other words, when learning the position of a goal in a maze surrounded by landmarks, children and older adults travelled, on average, a longer distance to the goal (travelled distance: F_(39,2)_=5.58, p<0.01, fig. 3b) and it took them longer to reach the goal (escape latency: F_(39,2)_=9.29, p<0.001, fig. 3c). To go further in the analysis, we separated the trial into an orientation period (which corresponds to the time it took the subject to start walking after the trial started) and a navigation period (in which the subject walk to the goal, see Methods). We found out that part of the longer escape latency was due to the fact that older adults took longer to start walking (orientation duration: F_(39,2)_=3.54, p<0.05, fig. 3d), likely indicating a lower confidence in taking decision when facing the environment composed of landmarks. Additionally, the average speed of the subject’s trajectory was lower (average speed: F_(39,2)_=4.69, p<0.05, fig. 3e, but also the normalized speed: F_(39,2)_=3.76, p<0.05, supp. fig. 4a) and thus the navigation period was longer (F_(39,2)_=9.22, p<0.001, supp. fig. 4b). Comparatively, the geometry version triggered no difference at all (travelled distance: F_(34,2)_=1.58, p=0.22, fig. 3b; escape latency: F_(34,2)_=1.47, p=0.25, fig. 3c; orientation duration: F_(34,2)_=0.71, p=0.5, fig. 3d; average speed: F_(34,2)_=0.32, p=0.73, fig. 3e; normalized speed: F_(34,2)_=0.87, p=0.43, supp. fig. 4a; navigation period: F_(34,2)_=1.79, p=0.18, supp. fig. 4b). Older adults and children thus travelled a similar distance, were as quick to start walking and to reach the goal zone than young adults, indicating a good level of confidence and efficient learning capacities when they are exposed to an environment with a geometric polarization. Additionally, the learning curves seem steeper in the geometry version, probably indicating a one-shot learning process in our participants.

We next wondered whether age influenced the way people explore the environment that could ultimately explain difficulty in using the landmarks. To do so, we estimated the intersections between the gaze vector and the virtual environment (see Methods) and we divide this data into 3 potentially informative areas of the environment, i.e. the sky region, the walls and the floor of the maze. Note that we did not record eye movements in children, hence analyses are restricted to the adult participants only. Figure 4 shows a double dissociation, with participants in the landmark version spending a higher proportion of time gazing at the sky regions (where the landmarks stand, U=986, p<0.0001), whereas people in the geometry version spend more time gazing at the floor region (U=856, p<0.0001, see supp. fig. 6 for data not averaged over learning trials). The average time spent on these critical areas of space was about 20% and people gazed to the maze walls for the remaining 80% of the trial time, independently of the versions considered (U=718, p=0.57). When visualizing spatial distribution of gaze intersections, we found out that people tend to look mainly at the circle landmark that was directly in front of the starting position in the landmark version and in the crotch area in the geometry version (heatmaps on fig. 4d,e). These critical areas were gazed during the beginning of the trial (mainly during the orientation period: supp. fig. 7). Importantly, older adults did not spend less time than young adults gazing to the sky region in the landmark version (U=119, p=0.30, fig. 4a), suggesting that the incapacity of older adults to use allocentric strategies relative to landmarks is not related to a default of attention to those landmarks during the learning process.

**Figure 3.**
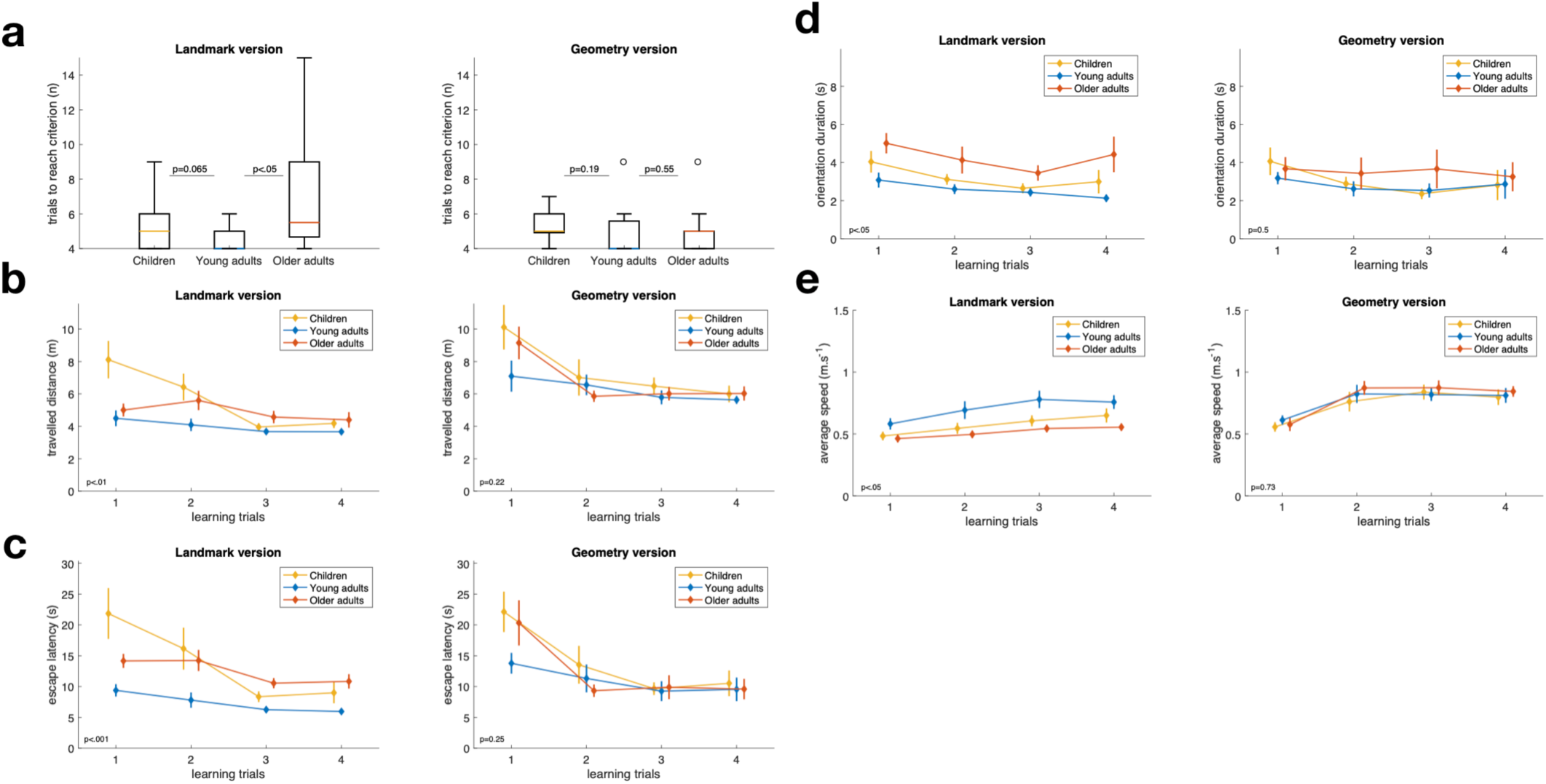
Spatial navigation performance of the three age groups during the learning phase, in the landmark version (left) and the geometry version (right). Overall, age differences are significant in the landmark version but not in the geometry version. P-value on the graphs corresponds to two-sample Wilcoxon rank sum test (a) or the main effect of age using one-way ANOVA for data averaged on the four first trial of the learning phase (b-e). Multiple comparisons of these data are provided in supp. table XX. Box plot representations show the median (coloured lines), the interquartile range (25th and 75th percentiles, length of the boxes), 1.5× interquartile range (whiskers) and outliers (circles, not present here). Position offset on the x-axis is added for clarity. Error bars show standard error.

**Figure 4.**
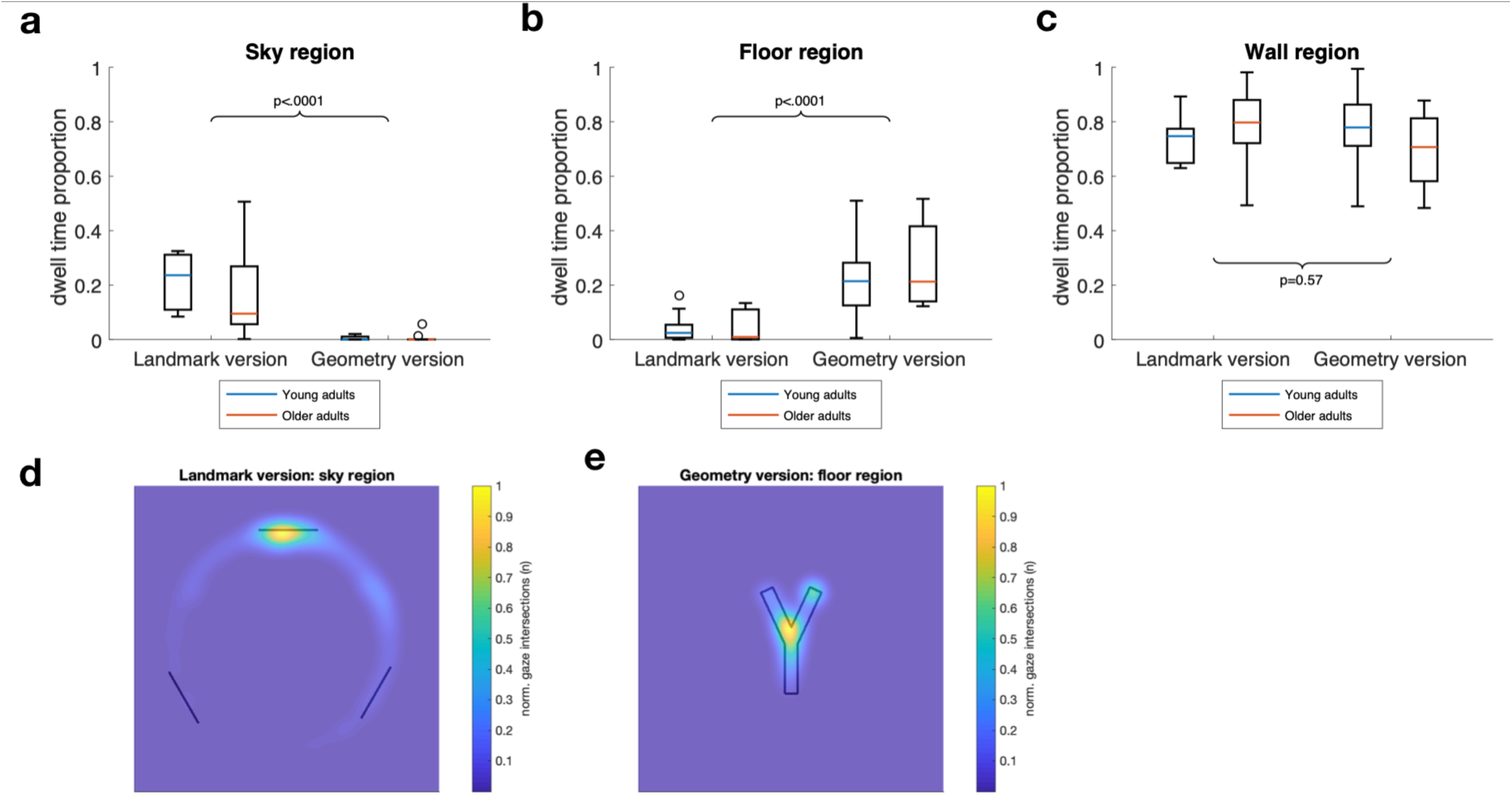
Dwell time proportion (a-c) and spatial distribution (d,e) of gaze for 3 areas of interest: sky, floor and wall regions. Participants in the landmark version spent significantly more time gazing to the sky region (particularly the circle landmark) whereas participants in the geometry version spent more time gazing to the floor (particularly the crotch region), independently of age. Data are averaged (a-c) or pooled (d,e) over the first four trials of the learning phase. Normality of dwell time proportion could not be reached, therefore P-values on graphs indicate non-parametric rank sum test comparing the two versions, with data are pooled on the factor age (a-c). Box plot representations (a-c) show the median (coloured lines), the interquartile range (25th and 75th percentiles, length of the boxes), 1.5× interquartile range (whiskers) and outliers (circles). Heatmaps (d,e) data are pooled across age and colorbar normalization is computed for each group separately.

To verify this point, we further separated our sample into egocentrers and allocentrers (see Methods on how we defined these two categories). Figure 5 shows that older egocentrers and allocentrers spent a similar proportion of time gazing at the sky region during the learning phase (t_(14)_=0.07, p=0.95). During the testing phase, however, we found an increased time spent gazing at the sky regions in people using an allocentric strategy (about 40% of the trial in both in young and older adults) whereas the egocentrers kept the same level as in the learning phase (20% of the trial, allocentrers vs. egocentrers in the older group: t_(14)_=2.24, p<0.05). To better describe this data, we further separated, among dwells directed to the sky, those directed to the circle, square, or star quadrants (fig. 5c) and estimated this variable over a window of 1s sliding over 15 (orientation period) and 35 (navigation period) time steps. Figure 5d shows that the young participants involved in an allocentric strategy gazed at the star quadrant (directly in front of the starting arm during the probe trials) during the beginning of the trial and this was apparently enough to understand where they were in maze and were the goal would be, indicating a good knowledge of the environment in this group. The older adults, although having the same early tendency to look at the star quadrant, exhibited an additional gaze component to the circle quadrant (fig. 5e), indicating that they might depend more heavily on the landmark experienced during the learning, that they might use it in a response-like manner (“to the right of the circle”). Finally, the egocentrers in the older group gazed at the star quadrant (about 30% of the time at the beginning of the trial), but apparently this does not elicit much in their decision (fig. 5f). Data in the geometry version shows that people using an allocentric strategy gazed at the floor region early during the probe trials, and no apparent difference between the behaviour in young or older adults was observed (supp. fig. 8). Given these clear differences, we next wondered whether we could predict the strategy chosen by the subject by observing its eye movements. We trained a binary classifier, on a single-subject-single-trial basis, with the altitude of the gaze, relative to the eye level, averaged during the orientation period of the probe trials (fig. 6ab) as a predictor variable. To assess the performance of the classified, we used a 25% hold out validation procedure on 1000 runs of the classifier and a leave-one-out validation (see Methods). We found that the model could predict which version of the maze the subject was assigned to by looking at the mean gaze altitude of the subject during the orientation period, i.e. when the subject did not even start to move in the maze (on 1000 runs: P_(performance<0.5)_ = 0.0001, leave-one-out: 88% of the subjects were correctly classified, n=49, fig. 6c). The spatial strategy chosen by the subject on probe trials could also be predicted, on a single-subject-single-trial basis, by gaze dynamics (on 1000 runs: P_(performance<0.5)_ = 0.05, leave-one-out: 79% of the subjects were correctly classified, n=26, fig. 6e), unlike the subject’s age which could not be predicted by gaze dynamic observation (on 1000 runs: P_(performance<0.5)_ = 0.5, leave-one-out: 55% of the subjects were correctly classified, n=49, fig. 6d).

**Figure 5.**
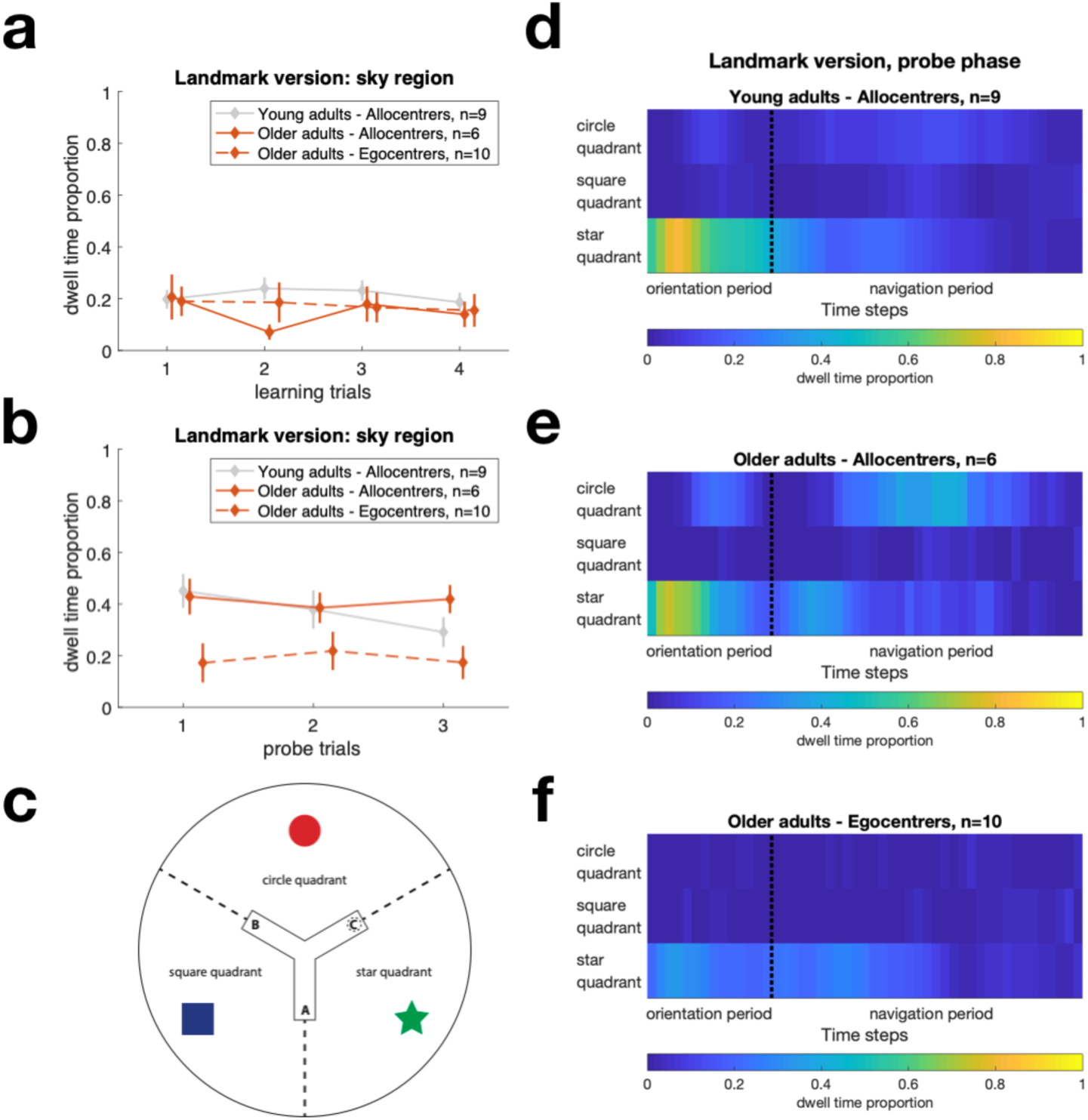
Gaze dynamics in the landmark version, separately for egocentrers and allocentrers. (a,b) Egocentrers did not spend a lower proportion gazing to the landmarks during the learning trials, however, they fail to increase this proportion during the probe trials. Position offset on the x-axis is added for clarity. Data from young allocentrers is shown for comparison. Data from young egocentrers is not shown, as there was only one subject using an egocentric strategy in this case. Error bars show standard error. (c) This proportion was further separated between the circle, square and star quadrants. (d) Young allocentrers looked at the star quadrant (which was in front of the starting arm) during the beginning of the trial (orientation period) and this seemed enough to understand where the actual goal was. (e) Older adults using an allocentric strategy had the same behaviour and show an additional component, gazing the circle quadrant (the one mostly experienced during the learning phase) during the orientation period and in the middle of the navigation period. (f) Older adults preferring the egocentric strategy still gazed at the star quadrant at the beginning of trial, indicating that the failure to use an allocentric strategy was not due to the fact that landmarks were not noticed.

**Figure 6.**
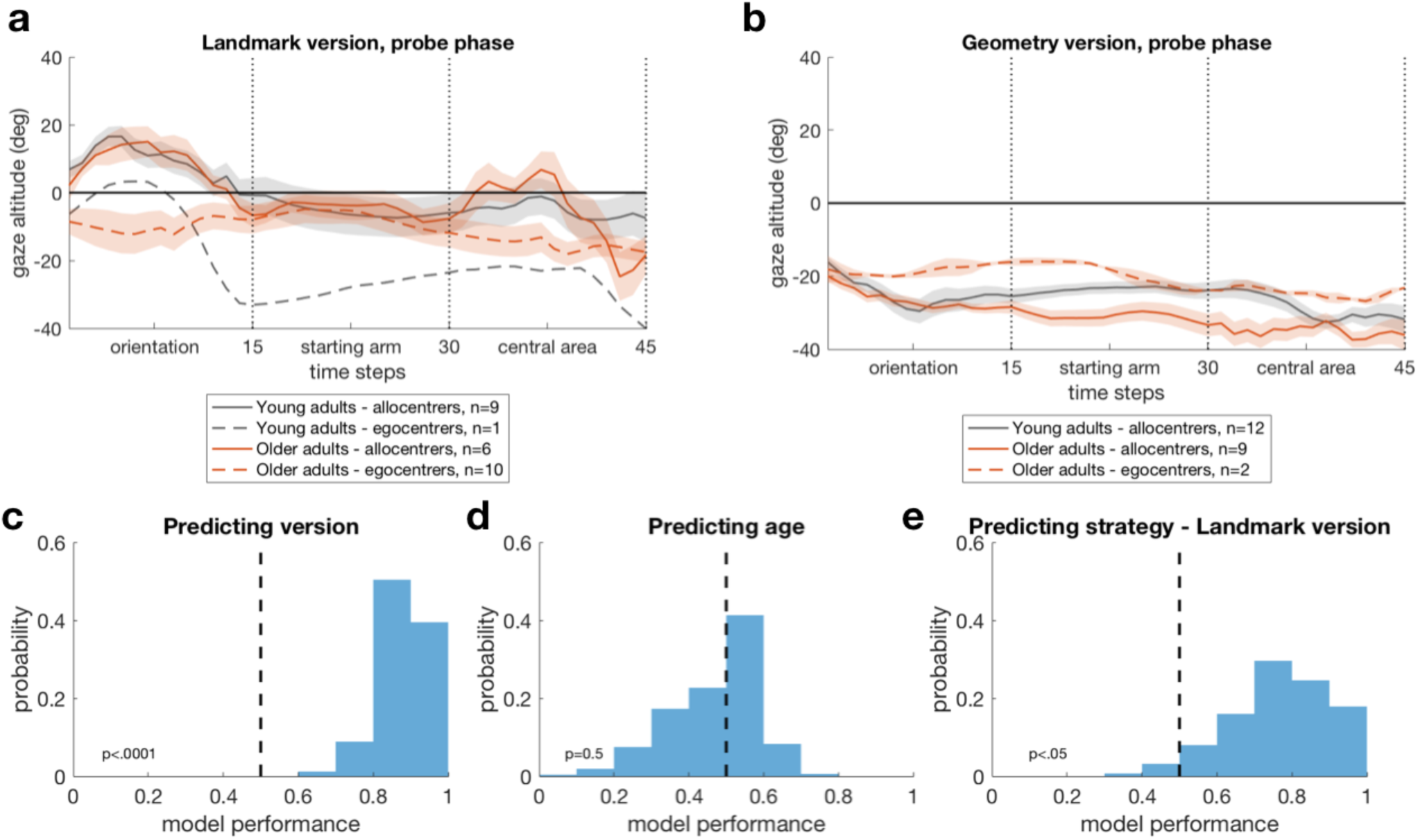
Prediction of the maze version, the subject age and strategy selection with eye movements. We used the altitude of the gaze, relative to the eye level, averaged during the orientation period of the probe trials (a and b) as a predictor variable in order to predict, on a single subject–single trial basis, the version of the maze performed by the subject (c), the subject’s age (d) and the strategy employed by the subject (e). The performance distribution of correct predictions on 1000 runs of the classifier suggests that gaze dynamics was predictive of the maze version and, for the landmark version, the strategy to be employed by the subject. The subject age could not be predicted with gaze dynamics. The indicated *P* value corresponds to P(performance < 0.5). Shaded areas on a and b indicate the between-subject s.e.m. The dashed lines on c, d and e indicate the chance level.

The additional gaze component to the circle quadrant in the older adults could indicate that older adults were not using the same process as young ones when employing an allocentric strategy. Indeed, although our paradigm was dedicated to separate the subject behaviour between allocentric and egocentric response, ecological navigation cannot be always accounted by such a clear dichotomy. In other words, a subject using allocentric navigation strategy can nevertheless employ various subprocesses of varying nature^20^. Figure 7 shows navigation variables, restricted to participants who employed an allocentric strategy on the probe trials. We found that using an allocentric strategy in relation to landmarks came at a time cost in older adults. Indeed, two-way analysis of variance showed a significant interaction between the factors age and version on time-related variables (normalized escape latency: F_(32,1)_=4.21, p<0.05, fig 7a; orientation duration: F_(32,1)_=6.95, p<0.05, fig. 7b; time in central area: F_(32,1)_=8.89, p<0.01, fig. 7c, but not on the distance-related variable, normalized travelled distance: F_(32,1)_=0.33, p=0.57, data not shown). Multiple comparison indicates that this interaction was due to older adults in the landmark version being longer to initiate walking, spending a longer time in the central area of the maze (up to 33 seconds in the first trial in one of our subjects, fig. 7c) and, as a result, being longer than young ones to reach the goal (supp. tab. 3). In comparison, in using an allocentric strategy in relation to the geometry, older adults were just as quick as young ones to make their decisions and as fast to reach the goal, arguing in favour of similar processes used in these two age groups (supp. tab. 3). The qualitative representations of the subjects’ behaviour during the first probe trial (fig. 7d,e) also supports the fact that older adults might not be using the same processes as young ones when employing an allocentric strategy, as it is defined by our experimental paradigm. Indeed, we observed that, unlike young adults, the older adults tended to stop in the central area of the maze and adopt the view experienced during the learning phase (i.e. gazing in the direction of the red circle), suggesting that these subjects might be trying to solve the probe trial with a view-matching behaviour rather than a purely allocentric one.

**Figure 7.**
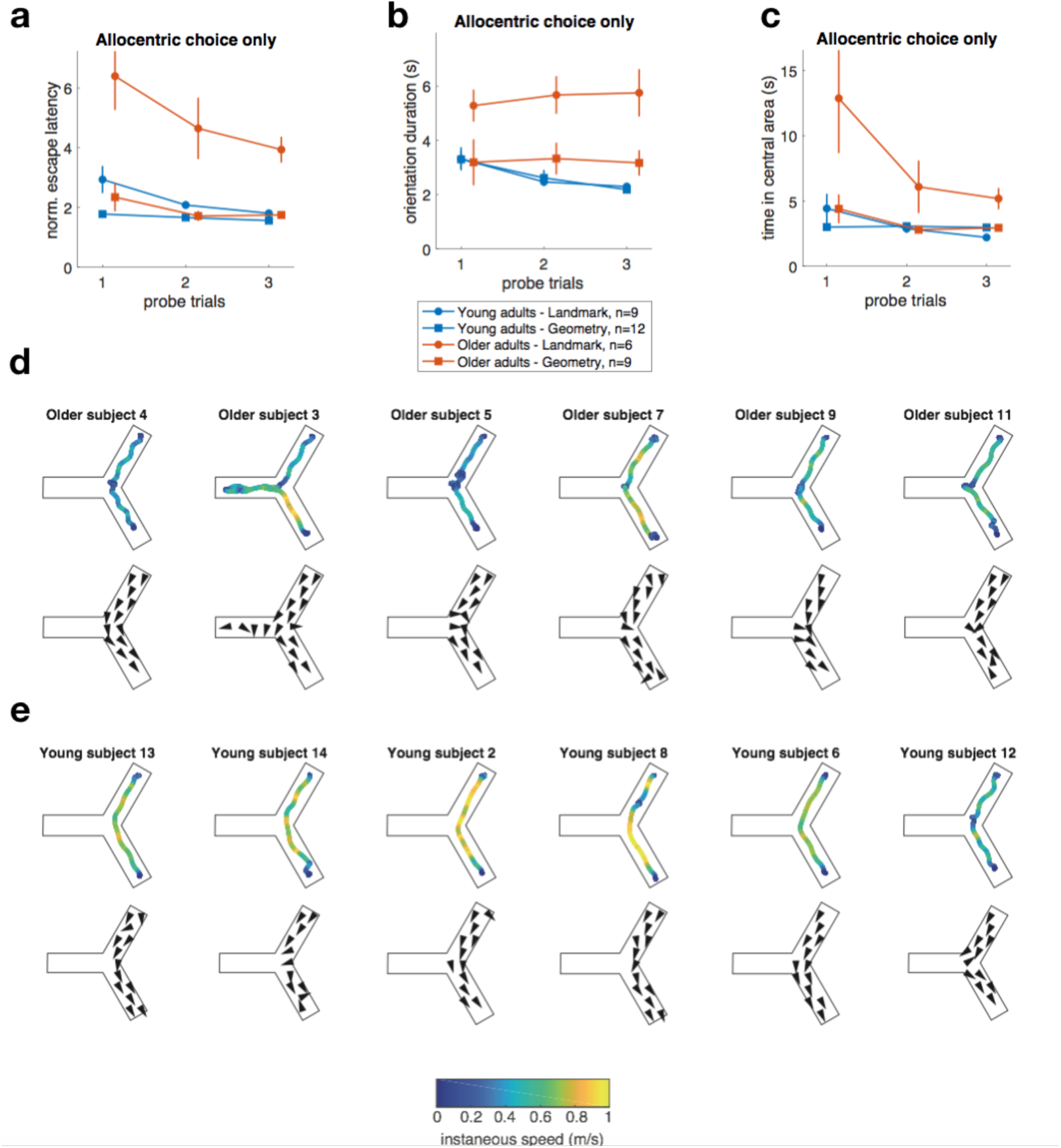
Different processes used by young and older adults while employing an allocentric strategy during the probe trials. (a-c) The observed time-related navigation variables suggest that, on one hand, using an allocentric strategy in relation to landmarks comes at a time cost in older adults. On the other hand, older adults are able to use geometric cues allocentrically with the same level of efficiency as young adults. Qualitative representation of the first probe trial in the 6 older adults (d) and 6 young adults (e) capable of using an allocentric strategy, as defined by our paradigm. The top row represents the trajectory and speed of the subject and the bottom row represents the gaze direction. Unlike young adults, older adults slowed down and even stopped in the central area of the maze. When they were in the central area, older adults were also more likely to adopt the view experienced during the learning phase (cf. arrow directed rightward), suggesting a view-matching process in these subjects. The escape latency (a) is normalized to account for the size difference between the two versions of the maze, the two other variables (b,c) can be directly compared between versions. Error bars show standard error.

We next wanted to know whether our subjects could be distinguished on visual or cognitive dimensions. For this analysis, we used the screening tests (see supp. tab. 4 for a detailed description of these tests) performed by some of our adult participants (up to n=48, depending on the test) in the Silversight cohort framework. We first used a principal component analysis (fig. 8a) in order to differentiate global visuo-cognitive profiles in our subjects. We found out that subjects could be discriminated on the first component based on their age (young vs. older participants: U=55, p<0.0001) but also, among the older groups, based on the strategy they used during the probe trials (allocentrers vs. egocentrers in the old group: U=117, p<0.01). When assessing the performance of subjects on each individual screening test, we found, here as well, a significant age effect on almost all tests (“Age effect” on fig. 8b). We also assessed, among the older group of subjects, whether visuo-cognitive functions differed between subjects who performed the landmark version or the geometry version (“Version effect”, on fig. 8b). This control was made to ensure that our sample was a priori homogeneous, and that an uneven distribution of visuo-cognitive profiles between the two versions of the maze could not explain our principal result, i.e. that older are better to use allocentric strategies in relation to geometric cues. We did not find any significant version effect in the older group (all p>0.1). Next, we wanted to compare profile of allocentrers and egocentrers, in the old group (“Choice effect” in fig. 8b). We found that egocentrers had lower capacity in terms of perspective taking (t_(24)_=2.63, p<0.05, fig. 8c), mental flexibility (U=167, p<0.05, fig. 8d) and contrast sensitivity, especially at high frequencies (fig. 8e, 0.5 circle per degree (CPD): t_(21)_=1.87, p=0.08; 1 CDP: t_(21)_=0.48, p=0.63; 2 CDP: t_(21)_=2.11, p<0.05; 4 CDP: t_(21)_=2.37, p<0.05; 8 CDP: t_(21)_=2.39, p<0.05; 16 CDP: U=101, p<0.001), when compared to allocentrers. Two-sample comparisons were also close to significance for the tests assessing 3d mental rotation (U=248, p=0.065) and working memory functions (U=249, p=0.0501, see complete data in supp. figs. 9 and 10). In other words, having a lower capacity to perceive fine details and lower executive and visuo-spatial abilities seems to be associated with a decrease usage of allocentric strategies.

**Figure 8.**
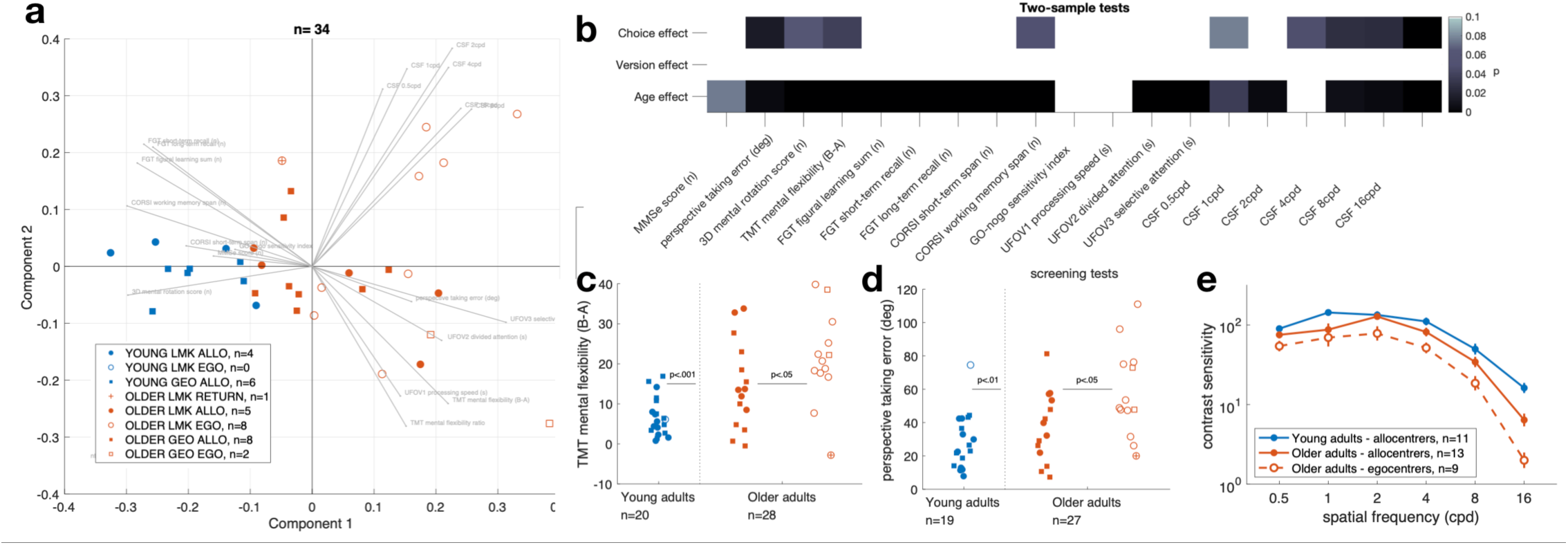
Visuo-cognitive profile of our participants. Among the older group, egocentrers had lower visual (particularly contrast sensitivity at high spatial frequency) and cognitive functions (particularly executive ones and visuo-spatial abilities), compared to allocentrers. Unsurprisingly, age also influenced almost all functions. (a) The principal component analysis combining all screening tests allowed us to discriminate profiles according to age and, in the old groups, according to the strategy used. (b) Overview of two-sample tests and the P-values obtained when comparing the age effect (young adults vs. older adults), the version effect (in the older group, geometry vs. landmark version) and the choice effect (in the older group, egocentrers vs. allocentrers). The P-values are uncorrected, only the darkest cells would pass correction for multiple comparisons. In the older group, egocentrers had lower mental flexibility (c), lower perspective taking capacity (d) and a lower sensitivity to contrast at highest spatial frequency (e). When the variable was continuous and normality of the data was reached, we used t-test. Otherwise, we used Wilcoxon rank sum test when the variable was ordinal. The size of the sample may differ between the analyses presented here. This is due to the fact that not all participants performed all the screening tests of the cohort. An offset is added to the x-axis for clarity (c,d). Error bars show standard error (e). MMSe: mini mental state examination; TMT: trail making test; UFOV: useful field of view; CPD: circles per degree.

Finally, a subset of subjects (n=20, 7 older adults and 13 children) performing the landmark version was asked, at the end of the experiment, to recognize the maze shape (among 3 possibilities, supp. fig. 11a), the landmarks (among 6 possibilities, supp. fig. 11b) and to draw a top-view map of maze they experienced. For the drawings, we assessed whether the constellation order was correct (supp. fig. 11c for a counterexample), the landmarks positioned correctly relative to the maze (supp. fig. 11d for a counterexample) and finally whether the goal zone was placed correctly relative to the landmarks. We found that the majority of subjects were capable to recognize the correct shape (85%) and the landmarks (100%) that they have seen during the experience. When drawing the map, however, approximately half of the subjects were not capable to remember the correct constellation order (45%), had a wrong landmark positioning (45%) and, as result, the goal positioning relative to landmarks was incorrect (65%, supp. fig. 11e). Interestingly, among the subjects who did at least one error on the drawing, 11 out 15 used an egocentric or return strategy in the testing phase, likely indicating that the landmarks are not bounded correctly to the representation of space, in those subjects.

## Discussion

By testing the navigation strategies used by subjects of three age groups in a maze composed of either landmarks or geometric cues, this study provides an evidence that geometric cues potentiate the use of allocentric strategies in children (∼10 yo) and older adults (>65 yo). When landmarks were the only source of information that could be used to define locations allocentrically, we observed a preference for egocentric strategies at both ends of the life course, forming an inverted-profile of strategies usage across the lifespan. When geometric cues could be used however, the pattern of results changed drastically. The proportion of spontaneous use of allocentric strategies significantly increased in both children and older adults, reaching the same level as young adults. By analysing in detail the behaviour of our subjects (head motion and gaze dynamics), we have shown that learning locations in an environment composed of landmarks solely was already more problematic in children and older adults, independently of the strategy preference. This age-related incapacity to use landmarks was not due to a default of attention, given that future egocentrers and allocentrers spent a similar proportion of time gazing at landmarks during the learning process. In the few older participants still capable to employ an allocentric strategy in relation to landmarks, our results indicate that the implementation of the strategy came at a large time cost, and that these subjects might be using one the three landmarks in a response manner (“to the right of”). We observed that egocentrers, although gazing at the landmarks, failed to use them to define locations. This was probably due to the fact that landmarks are not bounded correctly to the representation of space, early in development and with advancing age. We have found that providing geometric cues eliminates age differences in all the variables we observed: older adults and children were very efficient to learn locations in the environment geometrically polarized, and they were just as quick as young adults at making decisions and at implementing the strategy. This result indicates that older adults and children might not be as bad navigators as previously thought if geometric cues are available. Confirming previous results in real-world condition^18^, our results show that geometry is visually extracted by gazing at the floor, in a virtual environment as well.

Confronting our data with the existing literature allow us to confirm the preference for egocentric strategies in older adults, when the environment layout does not provide a geometric polarizing information and when performance, therefore, in based on landmarks only (using the Y-maze paradigm or others^7–12^). Regarding development, we complement previous evidence by showing a spontaneous preference for egocentric strategies in 10-year-old children when learning of a single stimulus-response association is required (as opposed to the more complex maze^6^). We add that this age-related strategy preference is not related to sensory restriction due to most-often used desktop virtual reality, given that our subjects show the same behaviour with an immersive head-mounted display, allowing both proprioceptive and vestibular information to be experienced during navigation. In addition, we proposed a divergent interpretation of this data, whereby the preference for an egocentric strategy observed in previous studies was actually conditioned by the sensory cues present in the environment (i.e. an issue with landmark processing) rather than a strategic choice per se. This interpretation fits well with other data showing that estimating and reproducing distances and rotations was relatively poor in older adults when the visual environment was composed of a circular (thus unpolarized) arena^21^ and that performance did not increase when landmarks were provided^22^. Our data fits also with paradigms of route learning, whereby retrieval of the contextual information (spatial position^23^ and/or temporal order^24,25^) about landmarks is impaired in older adults. The fact that free recall for landmarks was preserved in those studies is also confirmed by present results. In the light of the current study, we question whether path integration performance and route learning in large environments would be better in older adults and children when polarized layouts are used.

Altogether, our results suggest a critical function of geometry for orientation and navigation across the lifespan, which allows people of all age to quickly understand the layout, their own position within it and learn specific locations in space. Reference to geometry could thus represent a sort of default mode for spatial representation, which is well preserved across the lifespan, whereas landmark-based representation is developed at adult age. It may also require an additional cognitive effort for landmarks to be bounded to space representation. Our findings highlight the necessity to rethink the impact of age on spatial cognition and reframe the classical allocentric/egocentric dichotomy in order to integrate a landmark/geometry opposition that better explain age-dependent navigation deficits. It remains to be understood what makes geometry special relative to landmark and why we observe an age-related failure in anchoring the “cognitive map” with respect to landmarks. One possibility is that different sub-networks in the brain mediate the processing of geometric and landmark cues and that the sub-network dedicated to geometric processing is matured earlier in development and better preserved in aging. Experiments in our group are on-going in order to differentiate the brain areas implicated in geometry vs. landmark processing in humans, and characterize age-related cortical and sub-cortical dysfunctions potentially explaining why older adults and children are better at using allocentric strategies in the presence of geometric cues.

## Methods

### Participants

Seventy-nine subjects were included in this study: 29 children (range: 10-11 yrs, μ=10, δ=0.49, 17 females, 12 males), 22 young adults (range: 23-37 yrs, μ=28, δ=4.28, 13 females, 9 males) and 28 older adults (range: 67-81 yrs, μ=73, δ=3.90, 17 females, 11 males). The adult participants were part of the SilverSight cohort population (∼350 enrolled subjects) at the Vision Institute – Quinze-Vingts National Ophthalmology Centre, in Paris. The child participants were recruited in a primary school in the Paris area. All participants were voluntary and gave informed consent (parents gave informed consent for their child). The procedures were performed in accordance with the tenets of the Declaration of Helsinki, and they were approved by the Ethical Committee CPP Ile de France V (ID_RCB 2015-A01094-45, No. CPP: 16122 MSB). Adult participants were included in the study based on the following criteria: *i)* corrected visual acuity of at least 7/10, or 5/10, in participants younger or older than 70 years, respectively; *ii)* a Mini-Mental State Examination score of 24 or higher; *iii)* no physical inability in terms of locomoting without assistance (the complete list of inclusion/exclusion criteria used for the Silversight cohort are described in supp. tab. 5). The clinical and functional assessment of the Silversight cohort involved: ophthalmological screening (e.g., optical coherence tomography, fundus photography), functional visual screening (e.g., visual acuity, visual field extent, contrast sensitivity, attentional field of view), otorhinolaryngological examination (e.g., audiogram, vestibular function), cognitive-neuropsychological assessment (e.g., visuo-spatial memory, mental rotation, executive functions), oculomotor evaluation (e.g., ocular fixation, saccadic control), and a static/dynamic balance assessment. Among this multivariate assessment, we selected a subset of screening tests in order to control, as much as possible, for multiple co-factors at stake during spatial cognition, possibly entailing an unbiased interpretation of spatial behavioral data (e.g., with respect to inter-individual variability). These screening tests evaluated visual functions (contrast sensitivity at different spatial frequency) and cognitive functions (memory and executive functions, visuo-spatial abilities), which are detailed in supplementary figure 4. Participants habitually wearing far-vision lenses were encouraged to keep their glasses on during the experiment.

### Material

The experiment with the adult participants was performed in the Streetlab platform at the Institute of Vision and in a school gymnasium with the children participants. The virtual reality (VR) environment was created using the Unity3D game engine (Unity Technologies) and displayed in a HTC VIVE headset equipped with a Tobii Pro VR binocular eye tracker. Participants were equipped with a VR capable backpack computer (VR One, MSI). Experiment control and monitoring were performed remotely. This equipment allows the participant to move freely and explore the virtual environment with a feeling of immersion.

The real head position was tracked at 30 Hz by two laser emitters placed 9 m away form each other and at a height of 3 m allowing an experimental capture area of approximately 4.0 × 4.0 m. The HTC VIVE display had a nominal field of view of about 110° through two 1080 × 1200 pixels displays, updated at 90 Hz. The pixel density of the display was about 12 pixels/degree. The Tobii eye-tracker recorded eye movements at a rate of 120 Hz. The material used with the child participants was the same as for adults, with the difference that we did not use the Tobii eye-tracking integration with the children.

### Virtual environments

The two versions of the Y-maze were composed of 3 corridors, with walls covered by a non-informative homogeneous texture. The height of the walls was adapted to be 10 cm taller than the subject’s height, in order for all the subjects to have to exact same visual experience. In the landmark condition, the Y-maze had equiangular arms separated by 120° (fig. 1a). Each corridor was 66 cm large and 190 cm long. Three distal landmarks were placed outside of the maze, that is 8m above the walls and 20m from the centre of the maze. These were a green star, a blue square and a red circle, each subtending a visual angle of 10° relative to the centre of the maze. In the geometric condition, the geometric polarization of the maze was achieved by an anisotropic arrangement of the 3 arms, with the angle between arms being 155° for two sides and 50° for the last one. Each corridor was 66 cm large and 230 cm long and there were no distal landmarks in this condition. The corridors were longer in the geometric condition to prevent the subject from seeing the end of corridors when starting from any location. The maze was 1.54 times larger in the geometric condition with respect to the landmark condition. Subjects were randomly assigned to the landmark or geometry condition but we ensured an equal distribution of gender throughout the two groups. There were no shadows and the sky was homogeneous. Supplementary results 1 and supplementary figure 1(a-c) show that the performance of young adults between the geometry version and the landmark version were similar, arguing in favour of an equivalent level of difficulty in the two versions.

### Protocol

The experiment lasted 30min approximately and started with the calibration of the eye-tracking device. After adjusting the headset’s position and the inter-pupillary distance, the subjects performed a nine-points calibration without moving their head. To ensure the quality of the calibration procedure, a validation of the same nine points was performed. Whenever the mean angular error of the calibration was above 3°, the calibration process was started over. Validation (and recalibration if required) was performed at the beginning, halfway through and at the end of the experiment. During the experiment, the subject was disoriented before each trial. This procedure required the subject to hold the experimenter’s hands and be passively led around the room with eyes closed. A non-informative sound was display in headphones during the disorientation procedure to mask potentially uncontrollable sound from outside of the experimental room. We controlled that the disorientation procedure was truly effective by asking the subject to try pointing towards a computer, which was at the exit of the experimental room. Once disoriented, subjects were positioned at one of three starting positions (position A, B and C on Fig. 1), facing the center of the maze. At the end of each trial, the image displayed in the headset faded and the subject was instructed to close their eyes. Furthermore, the subject was told that walking through the virtual walls and standing on tiptoes were forbidden. The experiment proceeded as follows: during the “exploration phase”, the subject went through 3 exploration trials, of 60 seconds each, starting from one of the three starting positions (fig. 1, areas A, B and C). There was no specific task during these trials. The participant was instructed to explore the whole environment. Whenever the subject did not explore one of the corridors or did not look up in the direction of the landmarks, the experimenter gave a prompt by saying: “Make sure to explore the whole environment”. During the “learning phase”, the subject had to find a goal that triggered a rewarding sound. The goal was located at the end of corridor on the right (fig. 1, area C with dashed line which was 0.4m of radius). The starting location (fig. 1, area A) was the same throughout the learning phase. The subject was instructed to navigate as directly as possible to the goal zone. The learning phase ended after 4 consecutive successful trials, which were defined as a trajectory going directly to the goal zone without entering the corridor on the left. Then, during the “testing phase”, the subject was instructed to return to the goal zone and warned that there would be no rewarding signal this time. The starting position during these trials was changed, unknown to the subject, and followed the same pseudo-random order across subjects. The predefined order of starting location areas was: B, A, A, B, A, B. Trials starting from position B were the “probe trials”. Trials starting from A were the “control trials”, not analysed here. For the testing phase, the trial ends automatically when the subject stops for at least 5seconds in one of the three possible areas (A, B or C). At the end of the experiment, a subset of subjects (n=20) performing the landmark version was asked “subsidiary questions” by the experimenter. They were asked to *i)* indicate the shape of the maze they experienced among 3 different possibilities (supp. fig. 11a), *ii)* the landmarks that they noticed during the experiment between 6 possibilities (supp. fig. 11b), and *iii)* to draw a top-view map of maze they experienced (including walls, landmarks and goal position).

### Data processing

We first removed the two first seconds of recording, before which no image was displayed in the HMD. Then, we interpolated the head position (30 Hz) to fit the format of the eye-tracker data (120 Hz). The trial time was separated into an orientation period and navigation period. The orientation period starts when the image is displayed in the HMD until movement initiation, which is defined as the moment the subject surpasses a virtual circle of 0.3m around the starting position. The navigation period lasts until the subject enters the goal area (during the learning phase) or one of the three areas (A, B or C during the testing phase). Concerning the eye-tracking data, we estimated a cyclopean gaze vector by averaging data from the left and the right eye. If the signal from one eye was judged too noisy, by visual inspection, we used the data from the second eye. When then calculated the intersection of the gaze vector with 3 main environment plans: the walls, the floor and the sky region. For the later, we intersected the vector with a virtual sphere with a radius of 6m around the maze centre.

The navigation strategy was defined as the first area (A, B, C) entered by the subject on the probe trials. Whenever the subject first entered the actual goal location (fig. 1, area C with the dashed line), he/she was classified as using an allocentric strategy for that particular trial. Otherwise, if the subject represents the goal position relative to his own body position, he would navigate to area A and be using an egocentric strategy for that trial. Finally, the subject could also to return to the starting arm (B). This third, less observed possibility was termed a return strategy. We further separate our sample of subjects into “allocentrers” and “egocentrers” based on their choices on the probe trials. If a subject had a majority of allocentric or egocentric choices (2/3 or more), he/she was classified as allocentrer or egocentrer, respectively. Supplementary table 1 summarizes the number of observations and the mean and standard deviation of age in these two categories. Note that there were one children and one older subject who returned to the starting position on 2/3 or more probe trials (i.e. returning to area B) in the landmark version. Those two subjects are excluded from the analyses comparing allocentrers and egocentrers.

Several navigation variables were estimated, based on the trajectory of head positions, recorded by the HMD. Navigation performance was evaluated though several navigation variables. First, the number of trial needed to reach the learning criterion corresponded to the number of trials until reaching 4 consecutive trials where the subject goes directly to the goal area in the right arm, without entering the left arm (minimum: 4 trials). The escape latency and the travelled distance measured the time (in seconds) and distance (in meters) until reaching the goal zone. Given that the geometric version of the maze was slightly longer than the landmark one, we further normalized these two variables by the maze length. We also estimated the duration of the orientation period (from trial start until movement initiation, see definition above), the navigation period (from movement initiation until reaching the goal zone), and the time spent in the central area. The central area encompassed the last third of the three arms. Finally, we also estimated the average instantaneous speed of the trajectory from movement initiation until reaching the goal zone. Given that the walking speed is influenced by the subject’s height^26^ and that children were shorter than the adult participants, we also normalized the speed by the height.

The eye-tracking data were recorded for the adult participants only. The main variable we were interested to was the dwell time proportion, which corresponds to the proportion of time spent fixating either the walls, the floor or the sky region, normalized by the duration of the period considered. Missing data were not taken into account when normalizing this variable. There were, on average, a proportion of 0.27 and 0.22 missing data in the older group and the young group, respectively. Among the fixations directed to the sky, we further separated those directed to the circle, square, or star quadrant (fig. 5c). No particular attempt was made to separate fixations from saccades. Finally, for spatial distribution representations (i.e. heatmaps), we accumulated the gaze vector intersections and normalized the maps separately, for each group of subjects considered.

We used a generalized linear regression model in order to predict either the version of the maze the subject performs, his age or the spatial strategy he will employ on the probe trials. We used the averaged altitude of the gaze relative to eye level (in degrees) during the orientation period as an predictor variable and the version (i.e. landmark or geometry), the age (i.e. young or older) or the strategy (i.e. allocentric or egocentric) as binary response. We used two validation procedures: a 25% hold out validation on 1000 runs of the binary classifier where the trained model was tested on remaining 25% of the subjects, and leave-one-out validation, where the model was tested on the remaining subject. The performance of the model was assessed as the proportion or the number of correctly predicted response. The validation sets included all the three probe trials of a single subject, meaning that the model was not predicting some trials while being trained on the remaining trials of a single subject. The strategy was predicted for the subjects performing the landmark version only. Indeed, only two subjects choose the egocentric option in the geometry version, making it impossible for the model to learn appropriately.

Concerning the subsidiary questions, subjects were asked to recognize the maze shape among 3 possibilities (“shape recognition”, supp. fig. 11a,e), the landmarks that were present among 6 possibilities (“landmark recognition”, supp. fig. 11b,e) and to draw a top-view map of maze they experienced. For the scoring of the drawings, three other parameters were evaluated separately.

First, whether the landmark constellation was drawn in the correct order, whatever its position (“constellation order”, supp. fig. 11c,e). Second, whether the positioning of the landmarks was in-between arms and not at the end of the arm, whatever the constellation order (“landmark positioning”, supp. fig. 11e). Third, if the subject positioned the goal in the correct arm, relative to the landmarks (“goal positioning”, supp. fig. 11e).

### Statistical analyses

When the data were continuous and when normality and homoscedasticity allowed it, we used two-sample t-test (when comparing two subgroups), one-way ANOVA (when comparing three subgroups) and two-way ANOVA (when comparing two factors: age*version). Normality was verified by the Lilliefors normality test and visual inspection of Q-Q plots. Box-Cox transformation could be used to achieve normality and equalize variance^27^. We used two-sample Wilcoxon rank sum test or Fisher’s exact probability test for ordinal data and contingency table, respectively. Alpha level for statistical significance was set at *P* < 0.05.

## Acknowledgements

This research was supported by ANR—Essilor SilverSight Chair No. ANR-14-CHIN-0001. The funders had no role in the study design, data collection and analysis, decision to publish or preparation of the manuscript. We thank S. Mohand-Said of the Clinical Investigation Centre of the Quinze-Vingts Hospital, Paris, for medical supervision during clinical screening of participants. We also thank K. Lagrené, from the Aging in Vision and Action laboratory at Vision Institute, for helping in enrolling/profiling the participants. Finally, the authors wish to thank E. Gutman, J. Lebrun and C. Authié of the Streetlab team for technical support in setting up the experiments in the Streetlab platform.

## Author contributions

M.B., S.R., A.O.L., D.S., A.A. designed the experiment, M.B., A.O.L., S.R., G.T. collected and analysed the data, M.B., D.S., A.A. wrote the article.

## Supplementary information

### Supplementary results

#### Supplementary results 1

##### Comparing the level of difficulty of the two versions of the maze

We reasoned that if the landmark and the geometry versions of the maze had equivalent level of difficulty, we should observe no difference between the two versions in the basic measures of spatial learning in the group of reference, i.e. young adults. We thus considered the number of trials needed to reach the learning criterion (4 consecutive successful trials), the normalized travelled distance and escape latency (taking into account the fact that the two environments had different size) during the learning phase (supp. fig. XX). We found no significant difference in the performance of the young adults in the landmark in comparison to the geometry version on these three variables (trials to reach criterion: U=105.5, p=0.49; norm. travelled distance: U=118 p=0.87; norm. escape latency: t_(21,1)=_0.04, p=0.84), arguing in favour of a similar level of difficulty between the two versions, at least in young adults.

## Supplementary figures

**Supplementary figure 1.**
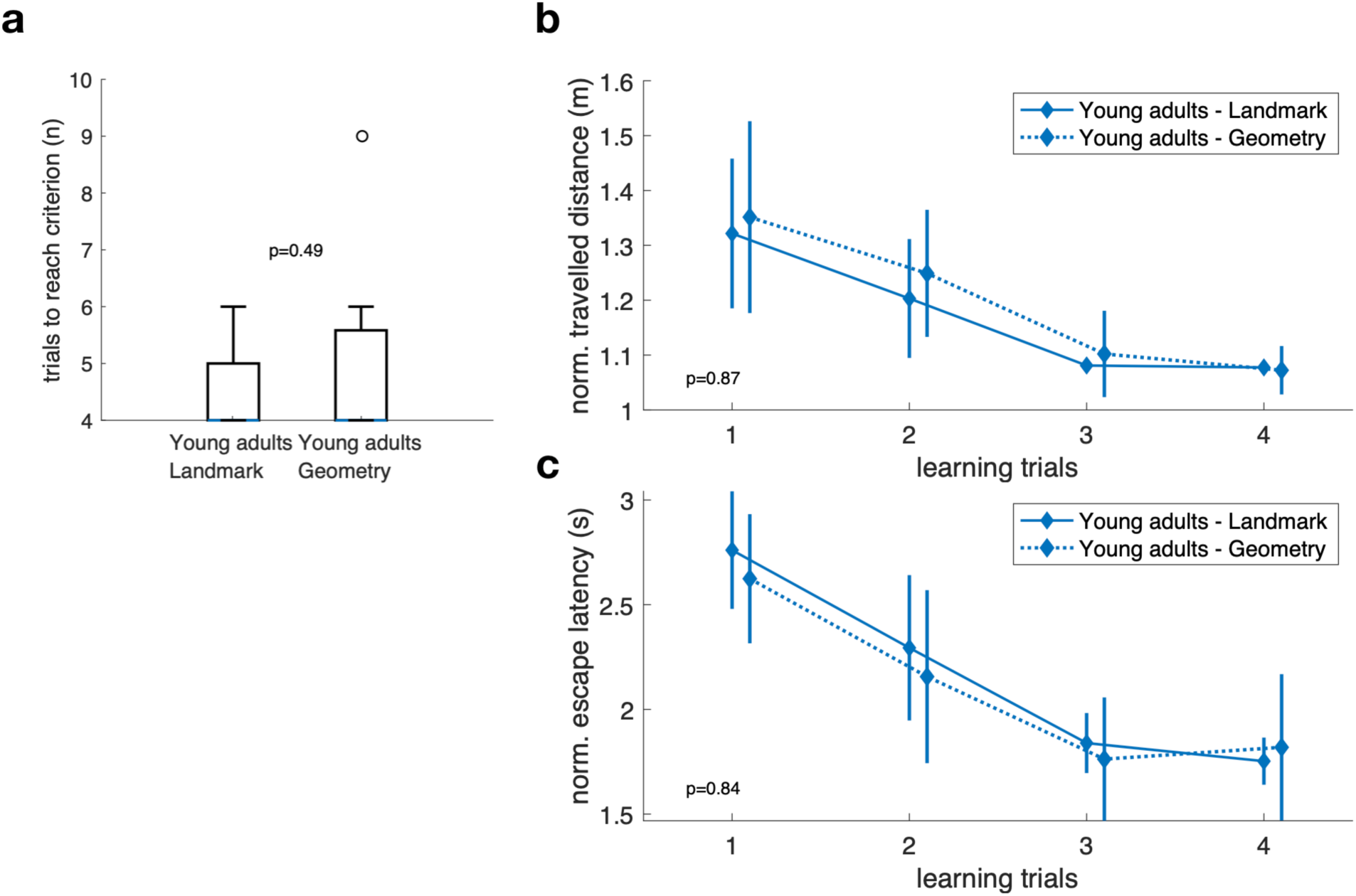
Comparing the level of difficulty of the two versions of the maze. There was no difference in the three basic measure of spatial learning between the two versions, in young adults, arguing in favour of an equivalent level of difficulty between the landmark and the geometry version. The travelled distance (b) and escape latency (c) are normalized to account for the size difference between the two versions of the maze, the number of trials needed to reach the learning criterion (a) can be directly compared between the versions. Data originates from the learning phase. Box plot representations showing the median (coloured lines), the interquartile range (25th and 75th percentiles, length of the boxes), 1.5× interquartile range (whiskers) and outliers (circles). The p-values on the graph indicate either parametric or non-parametric two-sample tests (*n* = 22). Error bars show standard error.

**Supplementary figure 2.**
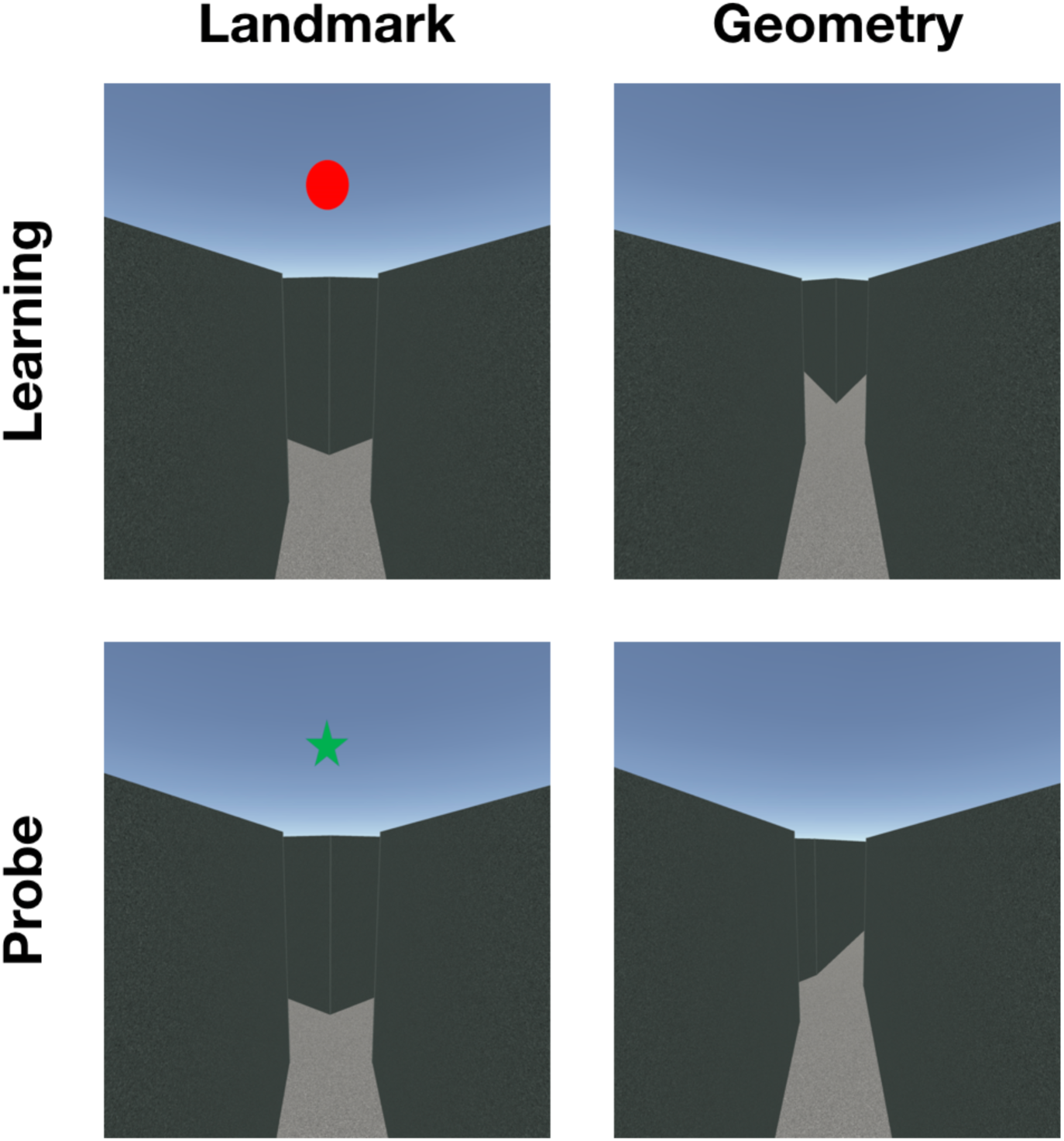
View experienced by the subjects at the beginning of a learning or probe trial, for the landmark and geometry version.

**Supplementary figure 3.**
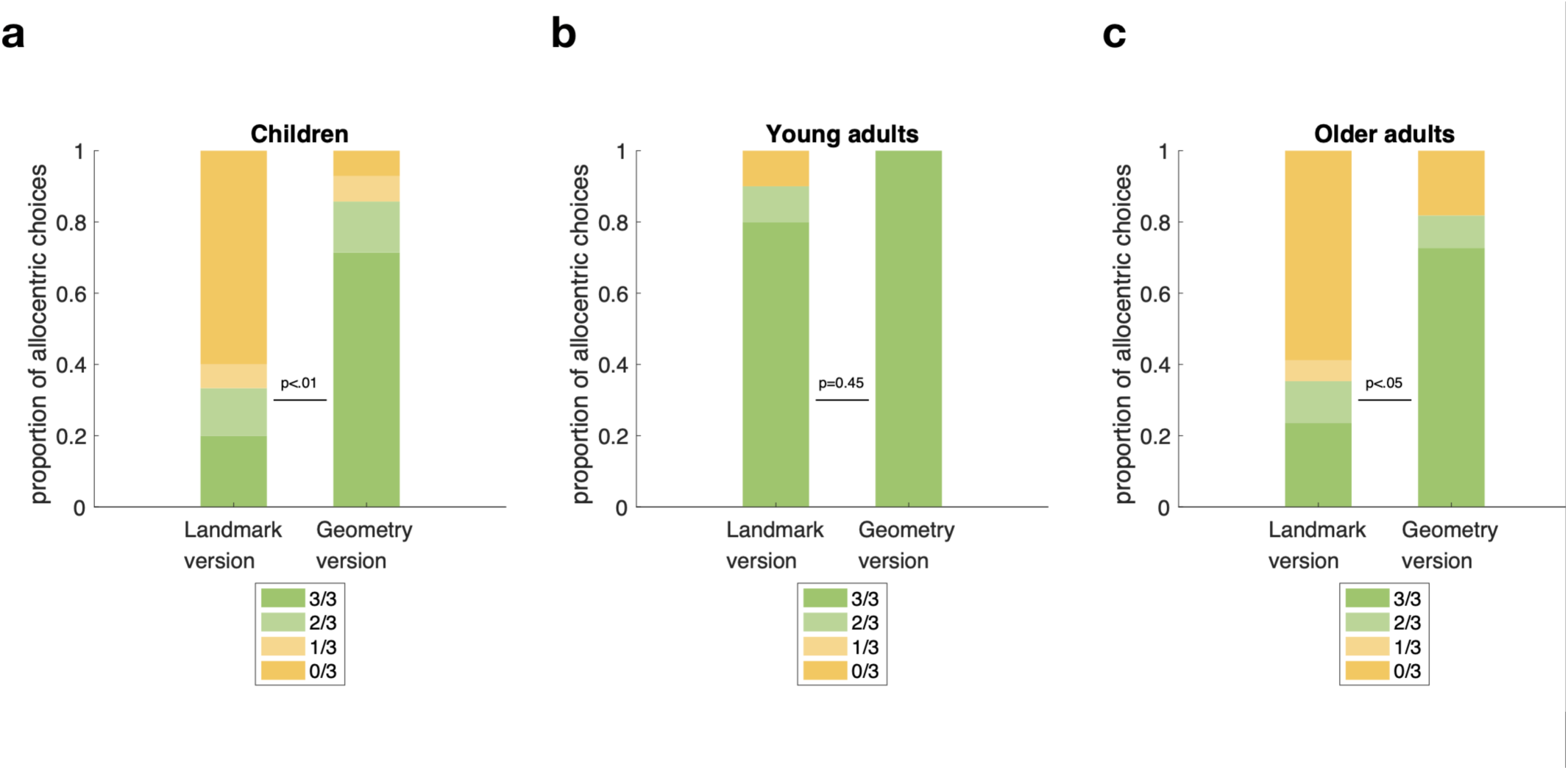
Proportion of allocentric choices for the three probe trials in the children (a), young adults (b) and older adults(c) for the two versions of the maze. In the children (n=29) and older adults (n=28), the proportion of participants using an allocentric navigation strategy was significantly lower in landmark version, in comparison to the geometry version. In young adults (n=22), there was no difference in the observed proportions between the two versions. P-values on the graph indicate Fisher’s exact test for two categories: majority of allocentric choices (green color: 3/3 and 2/3) and minority of allocentric choices (yellow color: 1/3 and 0/3).

**Supplementary figure 4.**
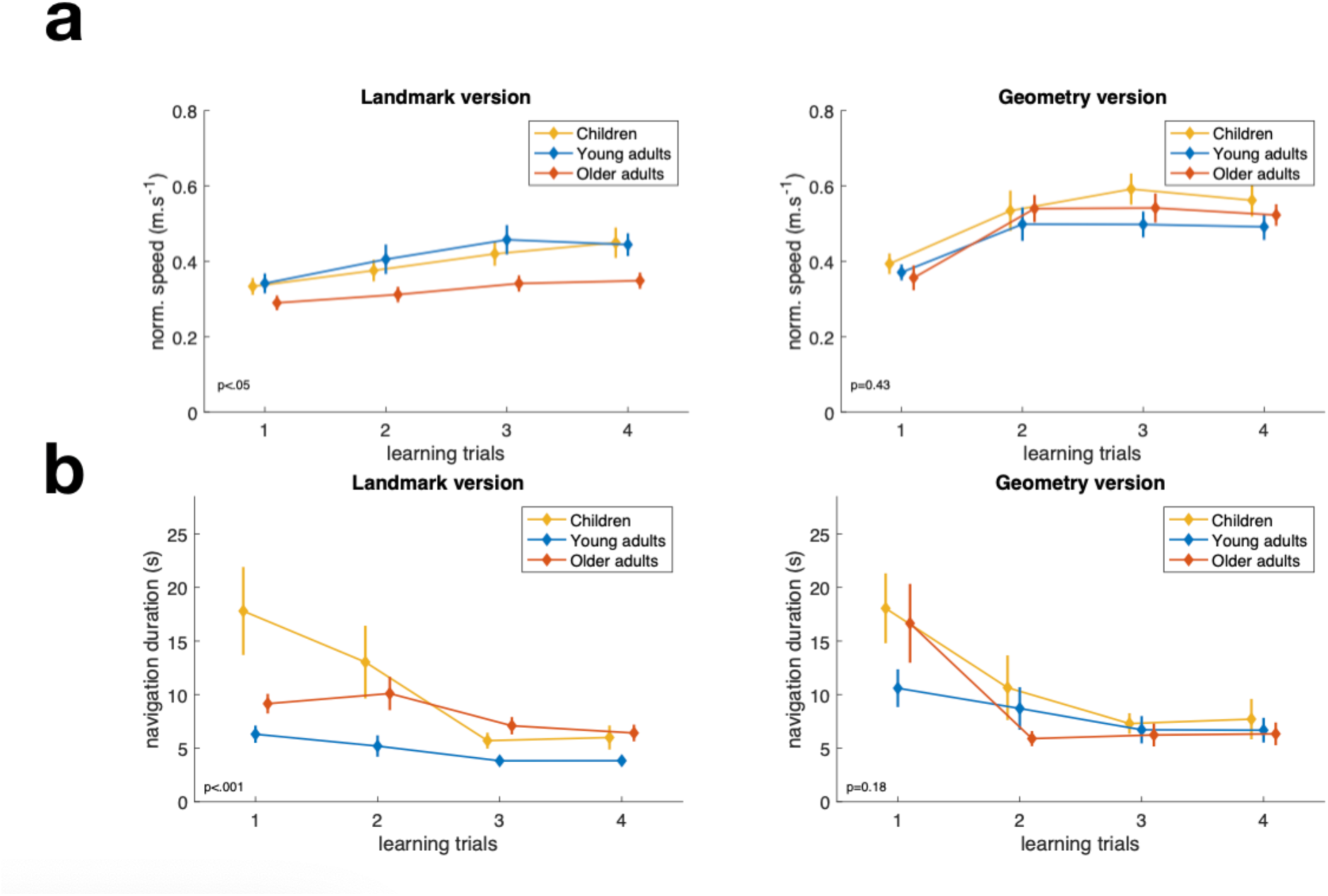
Normalized speed and (a) navigation period duration (b) during the learning phase for the landmark (left) and the geometry versions (right). Overall, age differences are significant in the landmark version but not in the geometry version. P-value on the graphs corresponds the main effect of age using one-way ANOVA for data averaged on the four first trial of the learning phase. Multiple comparisons of these data are provided in supp. table XX. Position offset on the x-axis is added for clarity. Error bars show standard error.

**Supplementary figure 5.**
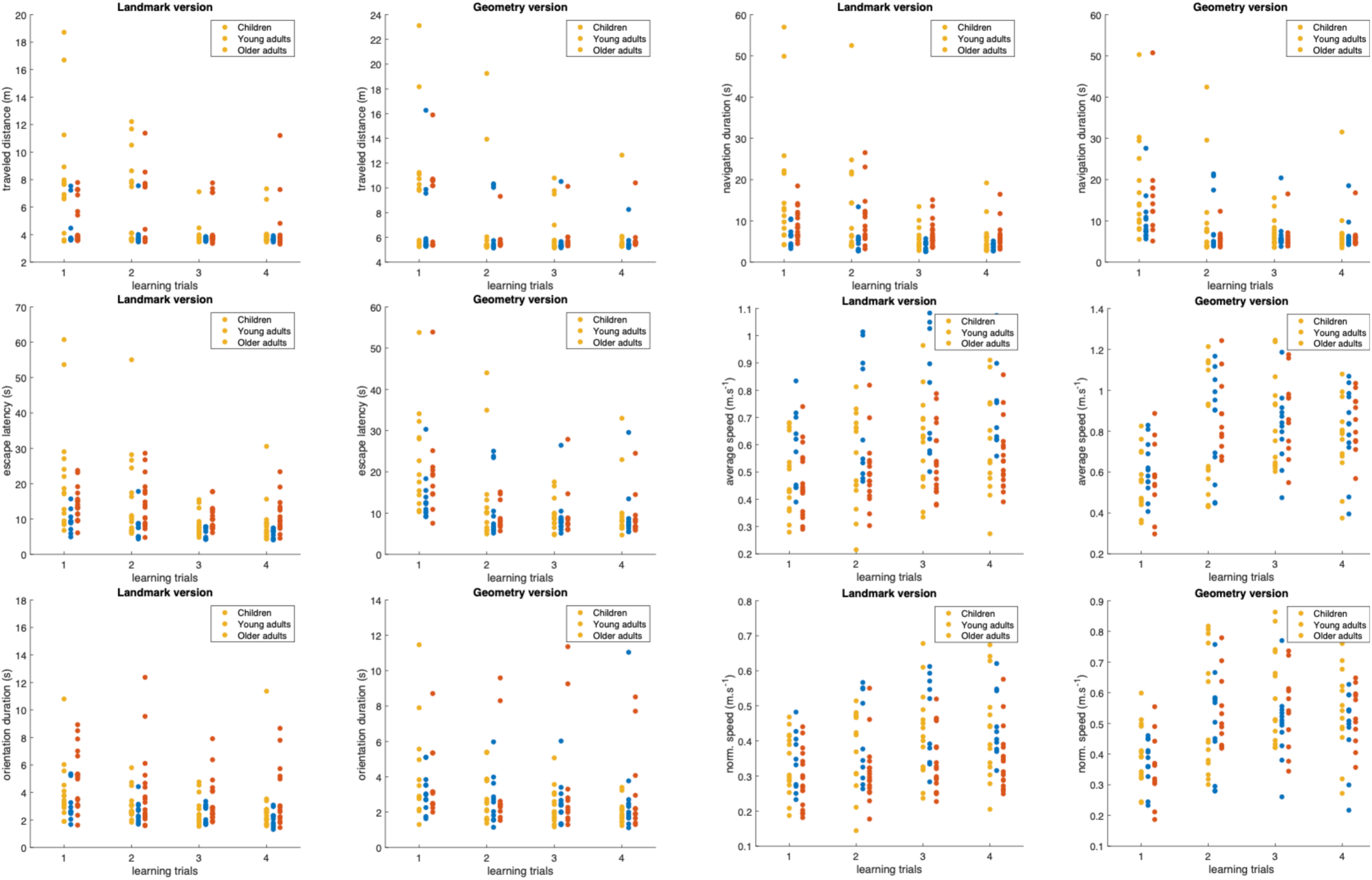
Scatter plots of the navigation variables for the first four trials of the learning phase, and the three age groups.

**Supplementary figure 6.**
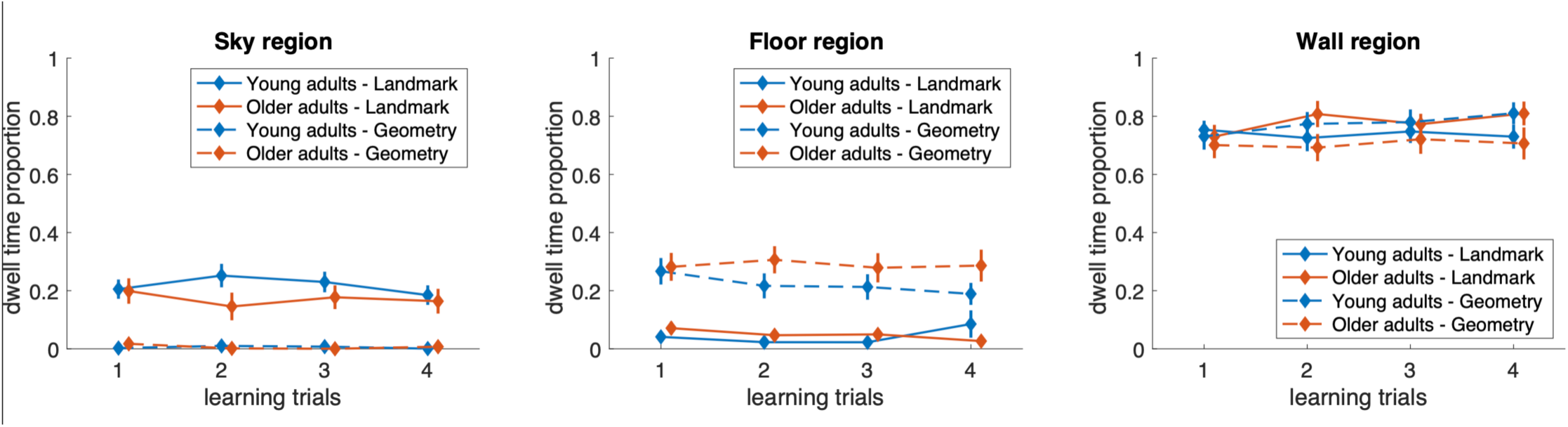
Dwell time proportion for 3 areas of interest: sky, floor and wall regions, for the four first trials of the learning phase. Participants in the landmark version spent significantly more time gazing to the sky region (particularly the circle landmark) whereas participants in the geometry version spent more time gazing to the floor (particularly the crotch region), independently of age. Position offset on the x-axis is added for clarity. Error bars show standard error.

**Supplementary figure 7.**
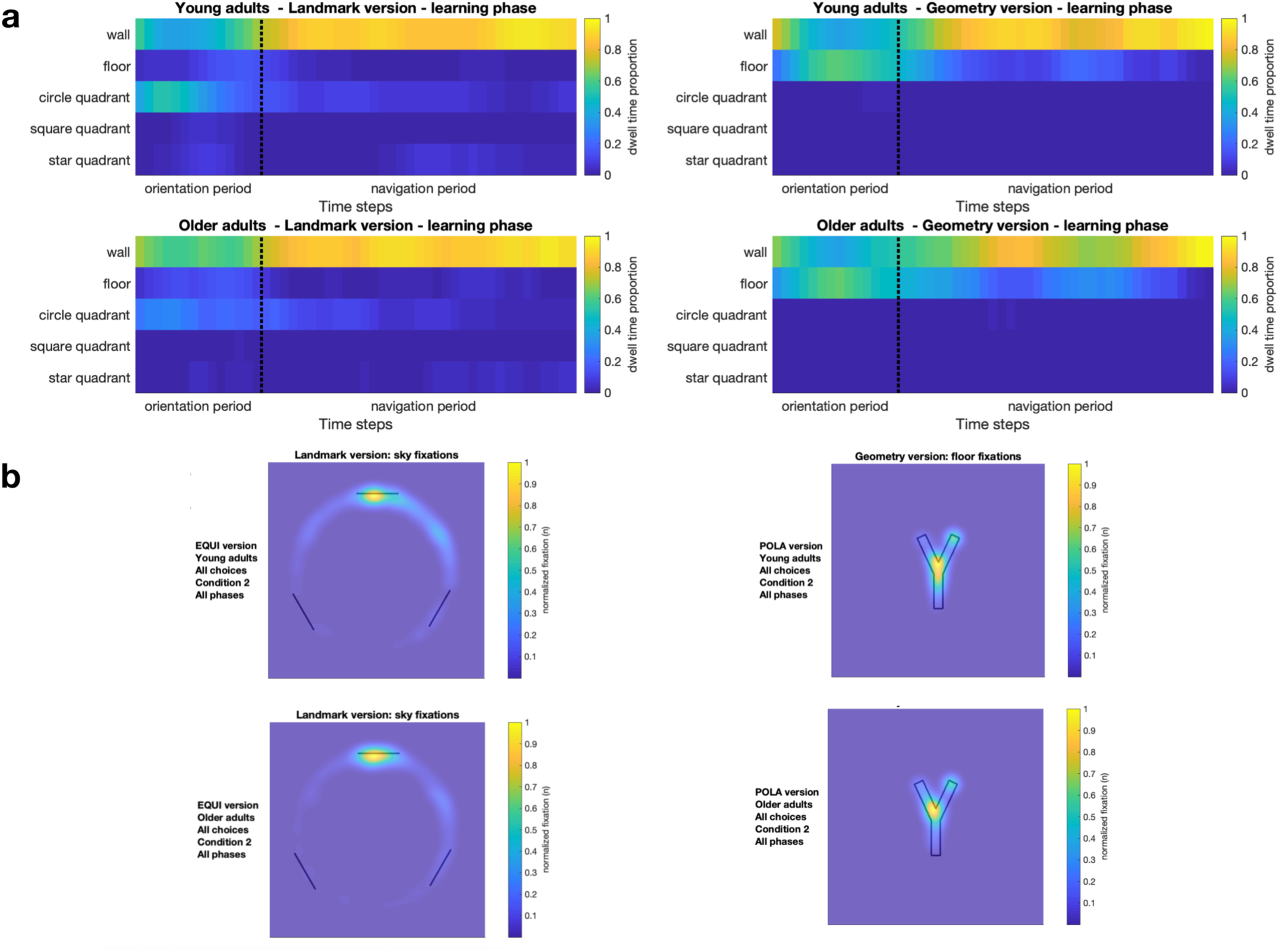
Dwell time proportion (a) and spatial distribution (b) of gaze for 3 areas of interest (sky, floor and wall regions) in young and older adults. Gaze dynamics did not differ between the two age groups. Data averaged (a) and pooled (b) across the 4 learning trials. Dwell time proportion was estimated over a window of XXs sliding over 15 (orientation period) and 35 (navigation period) time steps. Heatmaps (b) colorbar normalization is computed for each group separately.

**Supplementary figure 8.**
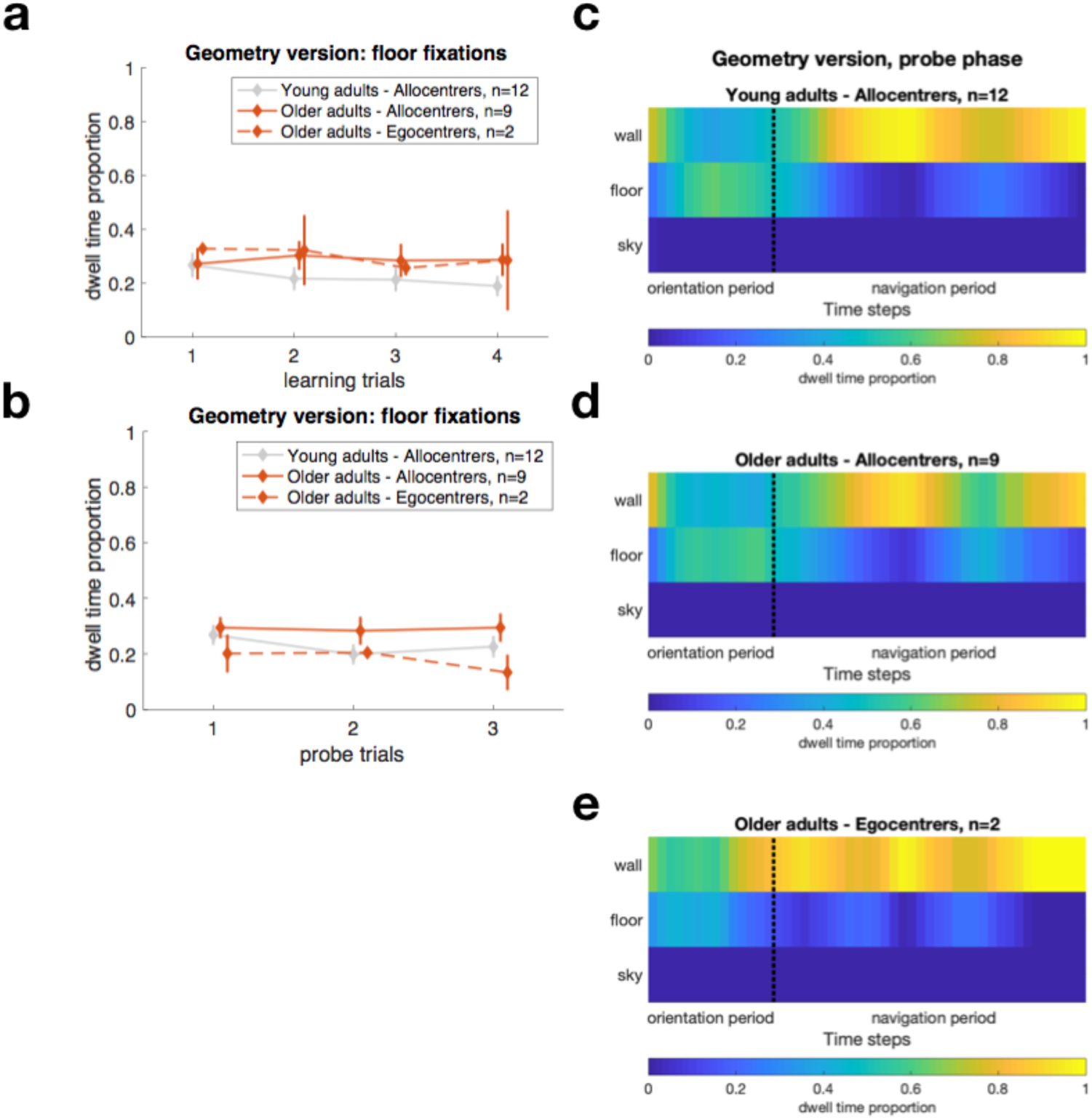
Gaze dynamics in the geometry version, separately for egocentrers and allocentrers. Dwell time proportion directed to the floor for the learning (a) and probe (b) trials. Data from young allocentrers is shown for comparison. Data from young egocentrers is not shown, as there was no subject using an egocentric strategy in this case. Position offset on the x-axis is added for clarity. Error bars show standard error. (c-e) Both young and older allocentrers had the same behaviour, gazing at the floor region at early during the probe trials. Comparison to the old egocentrers is difficult due to the small sample (n=2).

**Supplementary figure 9.**
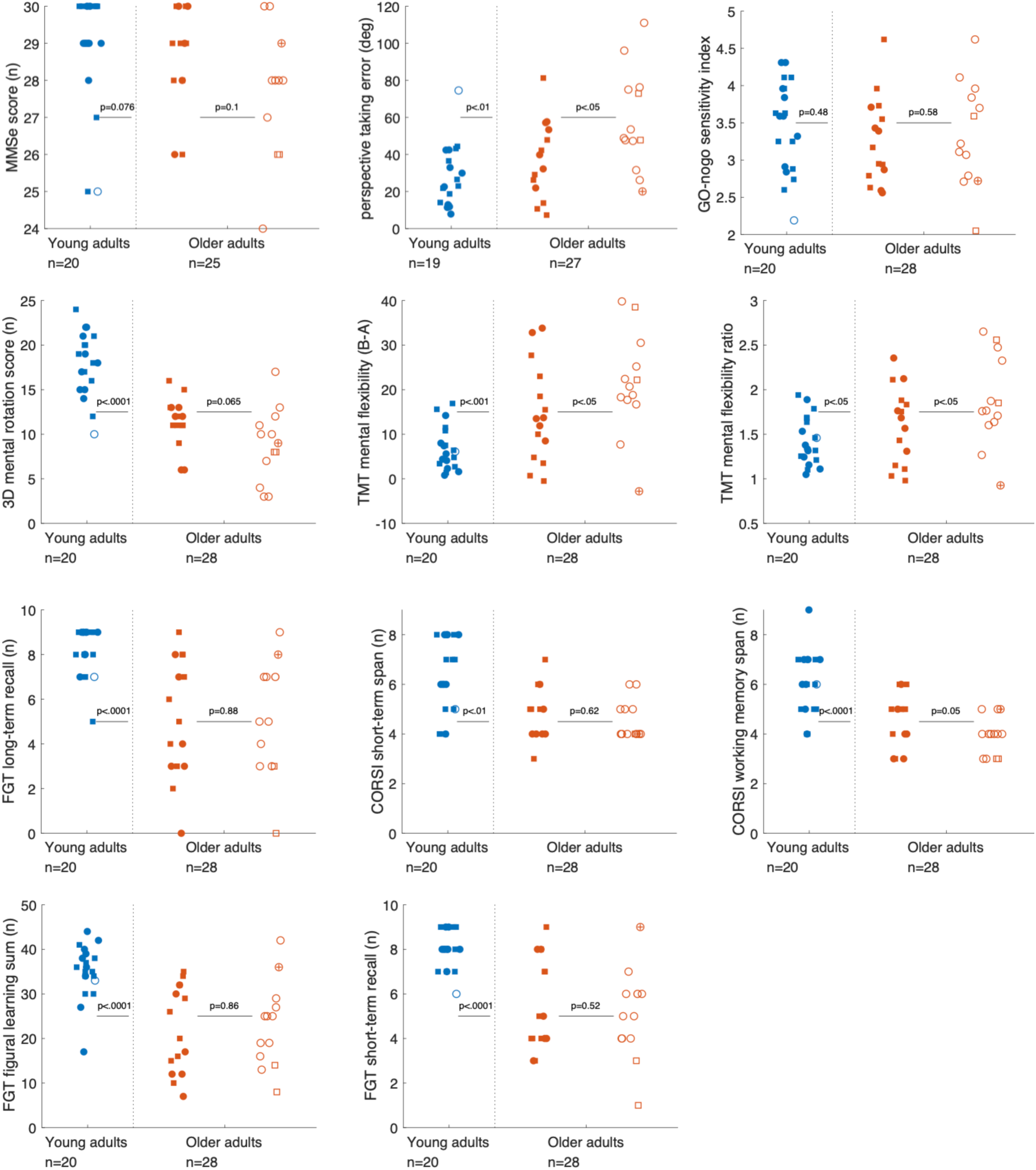
Cognitive functions in the adult participants. The P-value on the left side of the each corresponds to comparison of the “Age effect” (young vs. older adults, pooled across the two versions), whereas the P-value on the right indicates the “Choice effect” (allocentrers vs. egocentrers, in the old group). When the variable was continuous and normality of the data was reached, we used t-test. Otherwise, we used Wilcoxon rank sum test when the variable was ordinal. The size of the sample may differ between the analyses presented here. This is due to the fact that not all participants performed all the screening tests of the cohort. An offset is added to the x-axis for clarity. MMSe: mini mental state examination; TMT: trail-making test.

**Supplementary figure 10.**
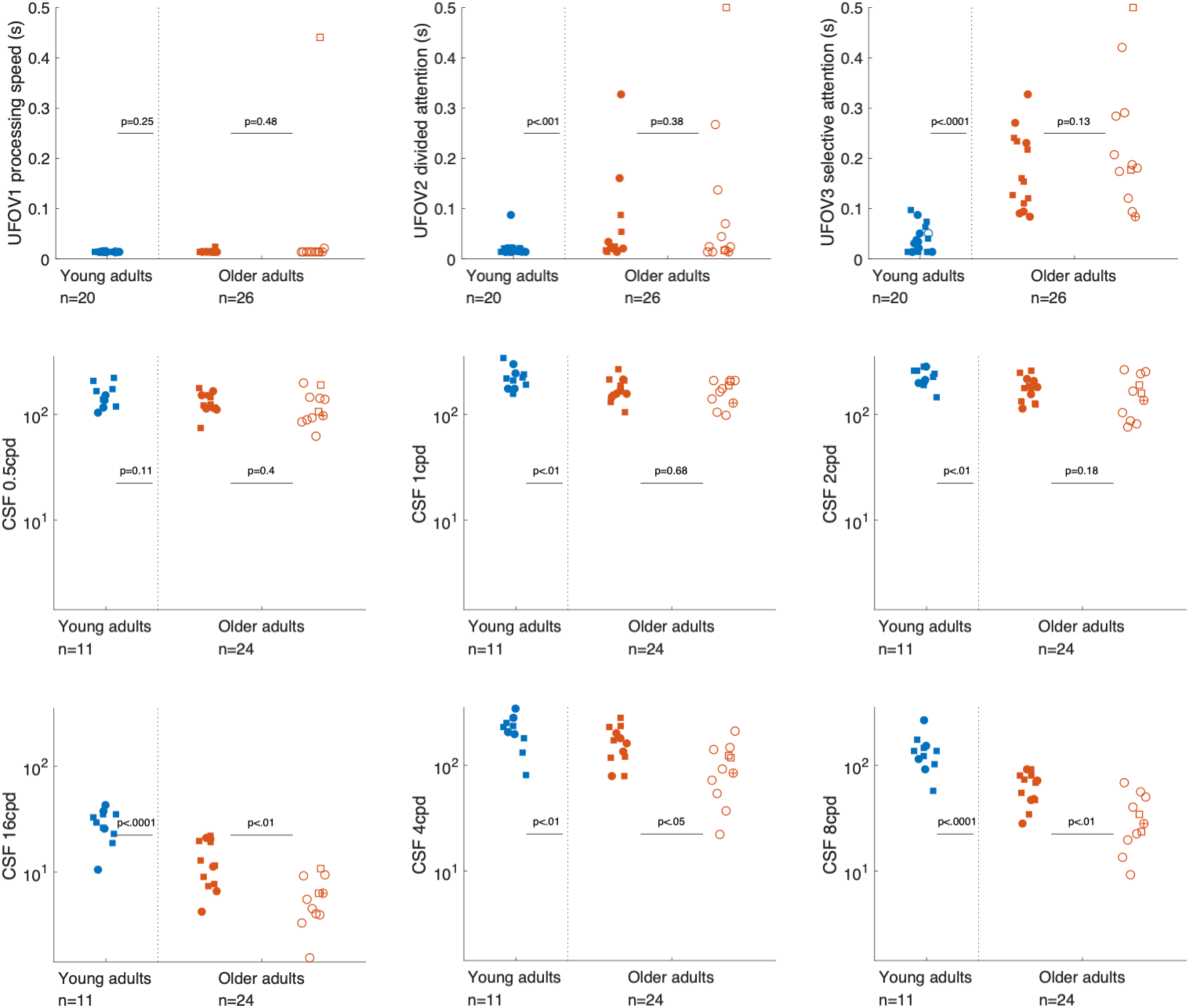
Visual functions in the adult participants. The P-value on the left side of the each corresponds to comparison of the “Age effect” (young vs. older adults, pooled across the two versions), whereas the P-value on the right indicates the “Choice effect” (allocentrers vs. egocentrers, in the old group). When the variable was continuous and normality of the data was reached, we used t-test. Otherwise, we used non-parametric Wilcoxon rank sum test when the variable was ordinal. The size of the sample may differ between the analyses presented here. This is due to the fact that not all participants performed all the screening tests of the cohort. An offset is added to the x-axis for clarity. UFOV: useful field of view; CSF: contrast sensitivity function; CPD: circles per degree.

**Supplementary figure 11.**
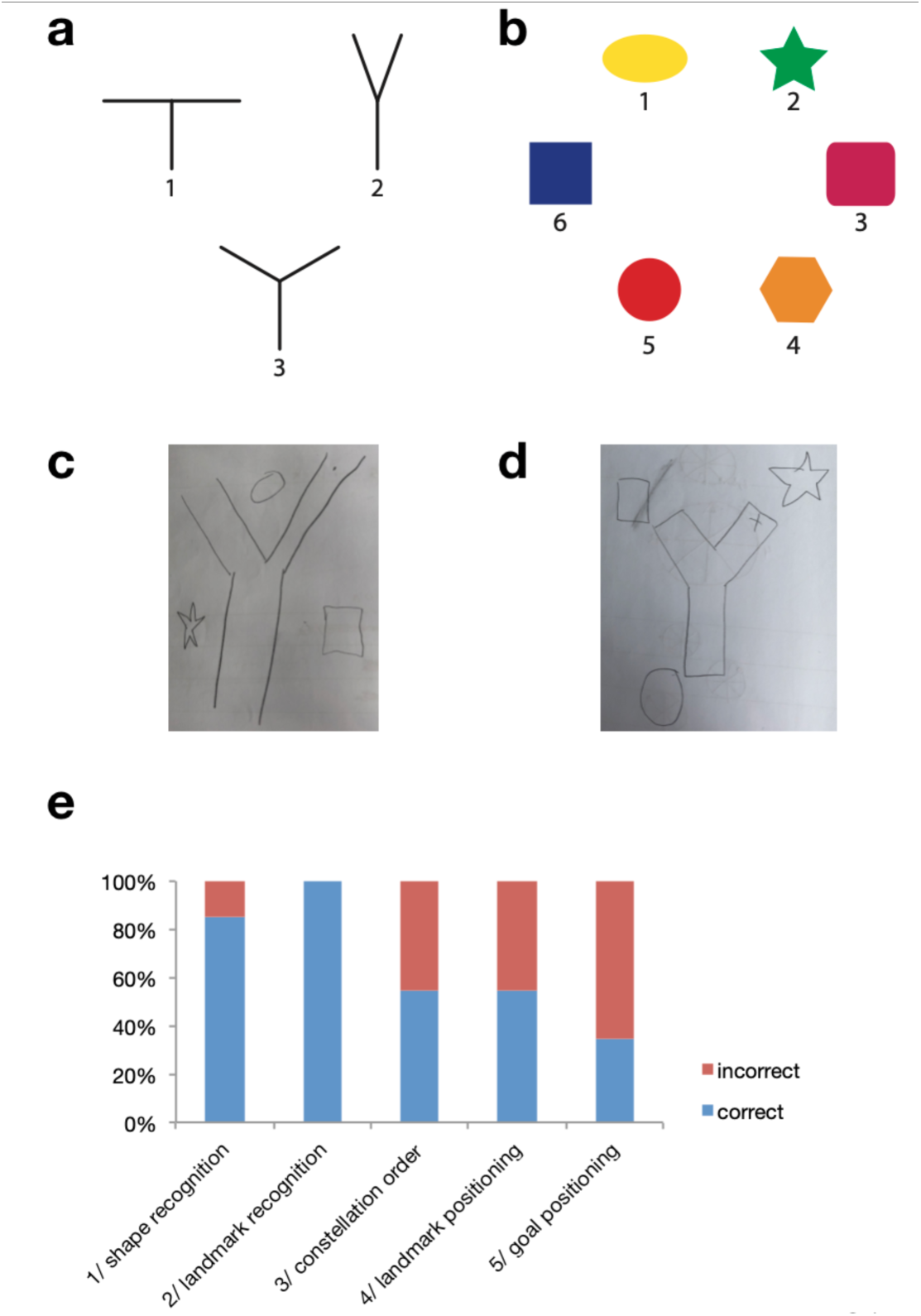
Subsidiary questions asked at the end of the experiment to a subset of subjects (7 older adults and 13 children) performing the landmark version. Subjects were asked to recognize the maze shape (among 3 possibilities, a), the landmarks that were present (among 6 possibilities, b) and to draw a top-view map of maze they experienced. (c) An example of drawing in which the landmark constellation order was incorrect. (b) An example of drawing in which the landmark positioning was incorrect (i.e. at the end of the arms instead of in-between). (e) Performance of the subjects on the 5 parameters assessed. Among the subjects that did at least one error on the drawing, 10 out 14 used an egocentric strategy in probe phase, likely indicating that the landmarks are not bounded correctly in the representation of space, in those subjects.

## Supplementary tables

**Supplementary table 1.** Number of observations, mean and standard deviation of age in subgroups. Allocentrers, egocentrers and returners are categories defined by the performance on the probe trials. For instance, a subject who had a majority of allocentric choices (2/3 or more) is categorized as an allocentrer.

|  |  | Children |  | Young adults |  | Older adults |  |
| --- | --- | --- | --- | --- | --- | --- | --- |
| | | n | mean age (± std) | n | mean age (± std) | n | mean age (± std) |
| Landmark version | allocenters | 5 | 10.40 (± 0.55) | 9 | 28.00 (± 5.35) | 6 | 72.67 (± 5.43) |
| | egocenters | 9 | 10.56 (± 0.53) | 1 | 25 | 10 | 73.00 (± 3.16) |
|  | returners | 1 | 10 | 0 |  | 1 | 76 |
| | total | 15 | 10.46 (± 0.52) | 10 | 27.7 (± 5.14) | 17 | 73.06 (± 3.93) |
| Geometry version | allocenters | 12 | 10.33 (± 0.49) | 12 | 28.08 (± 3.65) | 9 | 73.33 (± 4.53) |
| | egocenters | 2 | 10.00 (± 0) | 0 | | 2 | 72.50 (± 0.70) |
|  | returners | 0 |  | 0 |  | 0 |  |
| | total | 14 | 10.29 (± 0.47) | 12 | 28.08 (± 3.65) | 11 | 73.18 (± 4.07) |

**Supplementary table 2.**
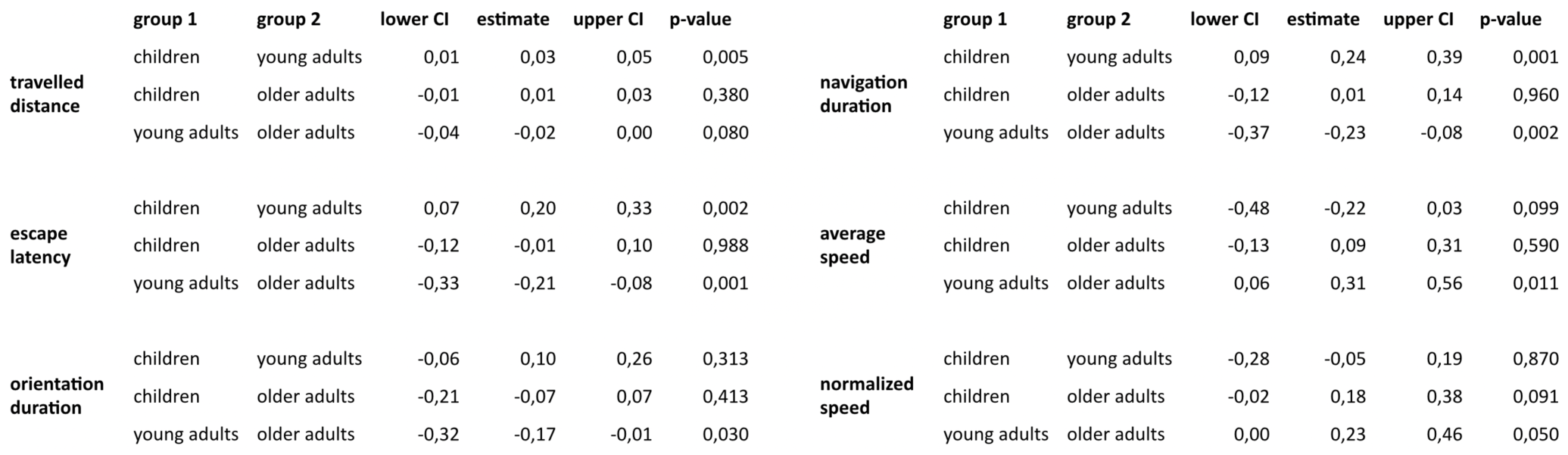
Multiple comparison of one-way ANOVA on navigation data, averaged over the first four trials of the learning phase. Tukey’s honest significant difference criterion is applied. CI: confidence interval of the estimate.

**Supplementary table 3.**
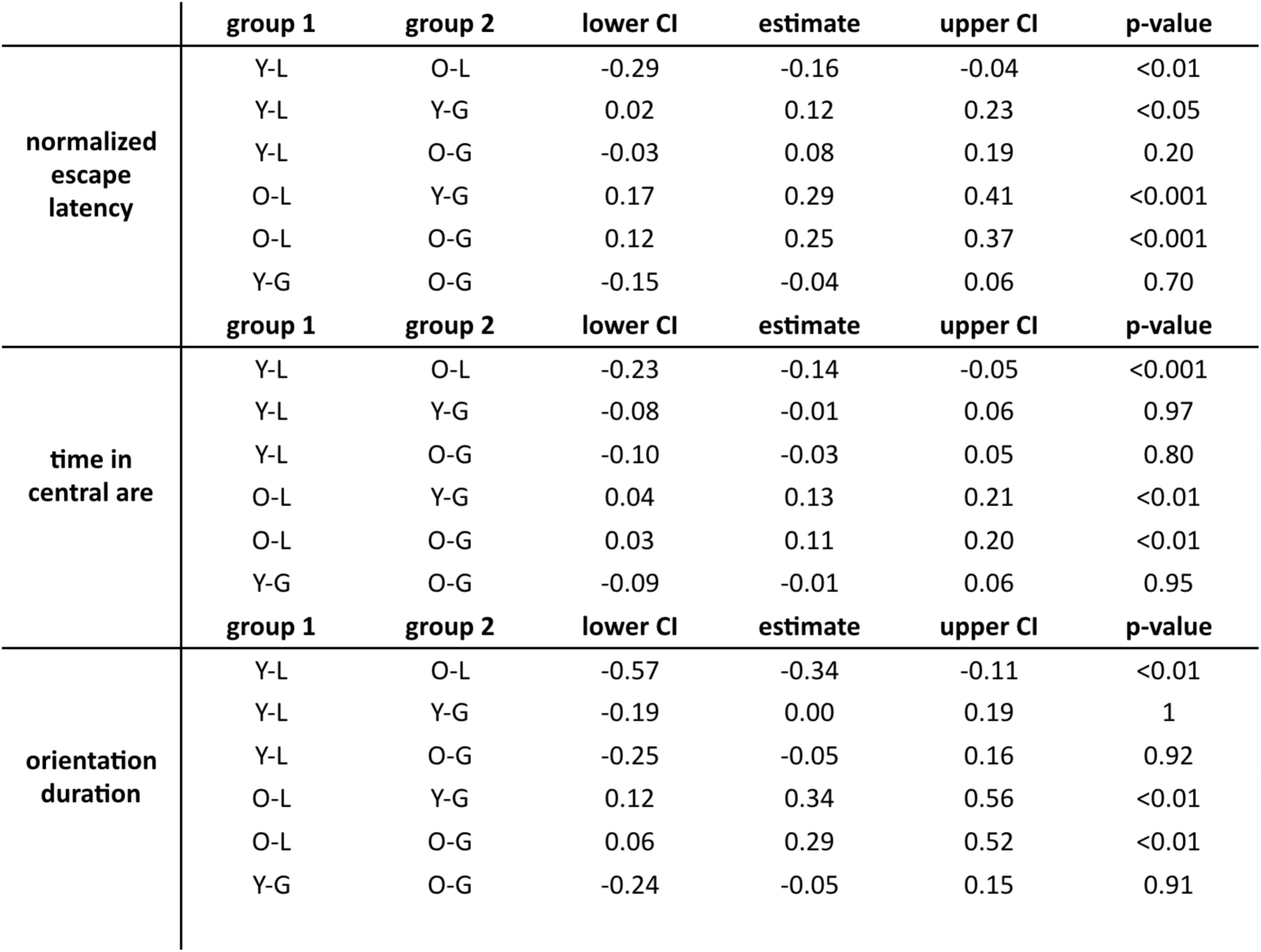
Multiple comparison of two-way ANOVA on time-related navigation variables, averaged over the probe trials. Tukey’s honest significant difference procedure is applied. CI : confidence interval of the estimate.

**Supplementary Table 4.** List of visual and cognitive tests performed by a subset of our adult participants.

|  |  |  |
| --- | --- | --- |
|  | Assessed function | Test/apparatus, task description and units of measurement |
| **Visual screening** | Contrast sensitivity | Evaluating sensitivity thresholds for different spatial frequencies (0.5, 1, 2, 4, 8, 16 circles per degree) in photopic condition, with subjects' own optical correction |
|  | Visual attention | Useful Field of View: central (UFOV1), central+peripheral visual discrimination task, without (UFOV2) and with (UFOV3) visual distractors, with subjects' own optical correction. Expressed in second needed for a correct discrimination (UFOV, Ball & Owsley, 1993) |
| **Cognitive screening** | Composite | Mini Mental State Examination (MMSE, Folstein, Folstein, & McHugh, 1975). Expressed as a score representing items succeeded. |
|  | Spatial working memory | Corsi block-tapping test: recalling spatial sequence of cubes in the forward (short-term span) or backward (working memory span) order (Schuhfried, 2004). Expressed as the largest sequence succeeded. |
|  | Visual memory | Figural Memory test: learning 9 visual figures presented 5 times and recalling them after 5 and 20 minutes (Schuhfried, 2004). Expressed as number of figures learned correctly (sum up to 45 items) or recalled (up to 9 items) |
|  | Mental rotation | 3D mental rotation test: mentally imaging views around 3D cubes (Schuhfried, 2004). Expressed as a number of correct items (up to 30) |
|  | Perspective taking | Perspective Taking/Spatial Orientation test: imagining position and facing direction relative to a two-dimensional array of objects and indicating the position of a third object (Hegarty & Waller, 2004). Expressed as an error in degree |
|  | Mental flexibility | Trail Making test: following sequences of letters (part A) and alternating between letters and numbers (B). Expressed as a difference between time needed in B and A (Schuhfried, 2004) |
| | Inhibition capacity | Go/no go task: responding to a frequent stimulus while inhibiting response to a rare one (Schuhfried, 2004). Expressed as a sensitivity index: $z(\text{hits}) - z(\text{false alarm})$ |

**Supplementary Table 5.** Inclusion/exclusion criteria used for the SilverSight cohort (adult participants only).

|  |  |
| --- | --- |
| **Inclusion criteria** | - Volunteer: man or woman<br>- Aged over 18 years old<br>- Affiliated to the social security system<br>- Absence of pathology, deficit or disorder that can interfere with visual, auditory, vestibular or cognitive functions |
| **Exclusion criteria** | - Person under guardianship<br>- Person using walker or wheelchair<br>- Person with a history of stroke<br>- Person with a history of epilepsy or convulsions<br>- Person with an history of active or progressive ophtalmological pathology to the exception of cataract<br>- Person with an history of active or progressive otological pathology or a surgical treatment (cholesteatoma, neuroma, otosclerosis)<br>- Best corrected visual acuity at 100% contrast lower than 7/10 or 5/10 before or after 70 yo, respectively, in one or both eyes<br>- Presence of abnormality on the monocular or binocular field of view<br>- Impaired color vision for one or both eyes on the D15 desaturated Lanthony test (protanope, deuteranope or tritanope)<br>- Ascending audiometric curve for one or both ears<br>- Balance disorders<br>- Minimal Mental State exam score lower than 24<br>- General Health Questionnaire score higher than 4 for depressive items |

## References

1. Burgess, N. Spatial cognition and the brain. Ann. N. Y. Acad. Sci. 1124, 77–97 (2008).

2. Ekstrom, A. D. & Isham, E. A. Human spatial navigation: representations across dimensions and scales. Curr. Opin. Behav. Sci. 17, 84–89 (2017).

3. Lester, A. W., Moffat, S. D., Wiener, J. M., Barnes, C. A. & Wolbers, T. The aging navigational system. Neuron 95, 1019–1035 (2017).

4. Newcombe, N. S. Navigation and the developing brain. (2019). doi:10.1242/jeb.186460

5. Nardini, M., Burgess, N. & Breckenridge, K. Differential developmental trajectories for egocentric, environmental and intrinsic frames of reference in spatial memory. 101, 153–172 (2006).

6. Bullens, J., Iglói, K., Berthoz, A., Postma, A. & Rondi-Reig, L. Developmental time course of the acquisition of sequential egocentric and allocentric navigation strategies. J. Exp. Child Psychol. 107, 337–350 (2010).

7. Colombo, D. et al. Egocentric and allocentric spatial reference frames in aging: A systematic review. Neurosci. Biobehav. Rev. 80, 605–621 (2017).

8. Wiener, J. M., de Condappa, O., Harris, M. a & Wolbers, T. Maladaptive bias for extrahippocampal navigation strategies in aging humans. J. Neurosci. 33, 6012–7 (2013).

9. Bohbot, V. D. et al. Virtual navigation strategies from childhood to senescence: evidence for changes across the life span. Front. Aging Neurosci. 4, 28 (2012).

10. Davis, R. L. & Weisbeck, C. Search Strategies Used by Older Adults in a Virtual Reality Place Learning Task. Gerontologist 55, S118–S127 (2015).

11. Driscoll, I., Hamilton, D. A., Yeo, R. A., Brooks, W. M. & Sutherland, R. J. Virtual navigation in humans: The impact of age, sex, and hormones on place learning. Horm. Behav. 47, 326–335 (2005).

12. Rodgers, M. K., Sindone, J. A. & Moffat, S. D. Effects of age on navigation strategy. Neurobiol. Aging 33, 997–1003 (2012).

13. Ruggiero, G., D’Errico, O. & Iachini, T. Development of egocentric and allocentric spatial representations from childhood to elderly age. Psychol. Res. 80, 259–272 (2016).

14. Harris, M. A., Wiener, J. M. & Wolbers, T. Aging specifically impairs switching to an allocentric navigational strategy. Front. Aging Neurosci. 4, 1–9 (2012).

15. Moffat, S. D. Aging and spatial navigation: what do we know and where do we go? Neuropsychol. Rev. 19, 478–89 (2009).

16. Lithfous, S., Dufour, A. & Després, O. Spatial navigation in normal aging and the prodromal stage of Alzheimer’s disease: insights from imaging and behavioral studies. Ageing Res. Rev. 12, 201–13 (2013).

17. Raz, N., Rodrigue, K. M., Head, D., Kennedy, K. M. & Acker, J. D. Differential aging of the medial temporal lobe: a study of a five-year change. Neurology 62, 433–438 (2004).

18. Bécu, M. et al. Age-related preference for geometric spatial cues during real-world navigation. Nat. Hum. Behav. (2019).

19. Hermer, L. & Spelke, E. A geometric process for spatial reorientation in young children. Nature (1994).

20. Ekstrom, A. D., Huffman, D. J. & Starrett, M. Interacting networks of brain regions underlie human spatial navigation : a review and novel synthesis of the literature. J. Neurophysiol. 118, 3328–3344 (2017).

21. Mahmood, O., Adamo, D., Briceno, E. & Moffat, S. D. Age differences in visual path integration. Behav. Brain Res. 205, 88–95 (2009).

22. Harris, M. a & Wolbers, T. Ageing effects on path integration and landmark navigation. Hippocampus 22, 1770–1780 (2012).

23. Jansen, P., Schmelter, A. & Heil, M. Spatial knowledge acquisition in younger and elderly adults: A study in a virtual environment. Exp. Psychol. 57, 54–60 (2010).

24. Head, D. & Isom, M. Age effects on wayfinding and route learning skills. Behav. Brain Res. 209, 49–58 (2010).

25. Wilkniss, S. M., Jones, M. G., Korol, D. L., Gold, P. E. & Manning, C. A. Age-related differences in an ecologically based study of route learning. Psychol. Aging 12, 372–375 (1997).

26. Samson, M. M. et al. Differences in gait parameters at a preferred walking speed in healthy subjects due to age, height and body weight. Aging Clin. Exp. Res. (2001).

27. Osborne, J. W. Improving your data transformations : Applying the Box-Cox transformation. Pract. Assessment, Res. Eval. 15, 1–9 (2010).

## Supplementary References

Ball, K. & Owsley, C. The useful field of view test: a new technique for evaluating age-related declines in visual function. J Am.Optom.Assoc. 64, 71–79 (1993).

Folstein, M. F., Folstein, S. E. & McHugh, P. R. Mini-Mental State: A practice method for grading the cognitive state of patients for the clinician. J Psychiatr Res 12, 189–198 (1975).

Schuhfried, G. Wienna test system (WTS). (2004).

Hegarty, M. & Waller, D. A dissociation between mental rotation and perspective-taking abilities. Intelligence 32, 175–191 (2004).

Tinetti, M. E., Richman, D. & Powell, L. Falls efficacy as a measure of fear of falling. J. Gerontol. 45, P239–P243 (1990).

Goldberg, D. P. & Hillier, V. F. A scaled version of the General Health Questionnaire. Psychol. Med. 9, 139–145 (1979).

